# Crusty-Pinky: A Novel Polyextremotolerant Fungus and its *Methylobacterium* Symbionts Could be an Essential Symbiosis for the Biological Soil Crust Consortium

**DOI:** 10.1101/2025.02.11.633654

**Authors:** Erin C Carr, Rajib Saha, Steven D Harris, Wayne R Riekhof

**Affiliations:** University of Nebraska-Lincoln; Iowa State University; University of Nebraska - Lincoln

## Abstract

JF2 08-2F Crusty is a novel melanized polyextremotolerant fungus isolated from a biological soil crust, which we believe harbors *Methylobacterium* spp. endosymbionts, called Pinky. Crusty is capable of utilizing many sources of carbon and nitrogen and is resistant to multiple metals and UV-C due to its melanized cell wall. We were unable to recover a Pinky-free culture of Crusty via usage of antibiotics. However, when exposed to antibiotics that kill or stop the growth of the Pinky, growth of Crusty is significantly stunted, implying that actively growing Pinky symbionts are needed for Crusty’s optimal growth. The Crusty-Pinky symbiosis also seems to be able to perform active metabolism in carbonless and nitrogenless medium, which we believe is due to Pinky’s ability to perform aerobic anoxygenic photosynthesis. Finally, Pinky was identified as being capable of growth stimulation of the algae *Chlorella sorokiniana*, indicating that Pinky likely produces cytokinins or auxins which *Methylobacterium* are known for. Features of this symbiosis provide us insight into the ecological roles of these microbes within the biological soil crust.

## Introduction

### Non-lichenized Fungi within Biological Soil Crusts

Biological soil crusts are dryland biofilms that form on the surface of non-vegetative mineral soils, primarily deserts, which rely on microbial interactions to survive the oligotrophic extremes of desert climates (Belnap et al., 2001; Belnap & Lange, 2003; Evans & Johansen, 1999). While the autotrophic members of the biological soil crust community (cyanobacteria, lichens, mosses, and algae) have relatively well established ecological niches, the niches of non-lichenized fungi and non-cyanobacteria bacteria are still yet to be discerned (Bates et al., 2006; Belnap, 2002; Belnap et al., 2001; Belnap & Lange, 2001; Csotonyi et al., 2010; Evans & Johansen, 1999; Harper & Marble, 1988; Johansen, 1993; Li et al., 2012; Maier et al., 2016; Steven et al., 2013). This is mostly due to the lack of research on these organisms within this ecosystem.

Deserts are notoriously harsh environments with extreme abiotic conditions such as intense UV exposure, diurnal temperature fluctuations, and osmotic pressures. Cold deserts often contain additional stressors such as light/dark annual extremes near the poles, drastic annual temperature changes, and excessive snow pack (Birkeland et al., 2020; Boulc’h et al., 2020). These extreme conditions create a difficult environment for active metabolism of the biological soil crust consortium, allowing for only short windows of opportunity with the optimal water availability and temperature conditions for active growth of the biological soil crust. Beyond that window, the community is in quiescence, and not actively respiring (Belnap, 2002; Belnap et al., 2004; Lange et al., 1997). Biological soil crust communities must cope with abiotic stressors simultaneously while relying on the autotrophic organisms to provide nutrients for the community during this short growth time period (Belnap & Büdel, 2016; Li et al., 2012). Given the abundance of such abiotic extremes, the biological soil crust needs to harbor multiple highly tolerant microbes to combat all causes of stress. This is because mechanisms underlying a given stress response are to some extent unique to that form of stress (ie: UV exposure vs. water retention) (Bowker et al., 2002). Additionally, the sources of abiotic stress resistances are not well understood, partly because the microbes present within the biological soil crust are not very well characterized. One source of abiotic resistance that multiple microbes in the biological soil crust are known to possess is sunscreen-like compounds and other UV-resistant mechanisms such as carotenoids or melanin (Bowker et al., 2002; Gao & Garcia-Pichel, 2011).

Polyextremotolerant fungi are a paraphyletic group of fungi characterized by their melanized cell wall which has imbued them with high abiotic tolerance (Gostinčar et al., 2009; Gostinčar et al., 2012). These fungi have previously been isolated from biological soil crusts (Bates et al., 2006), but their ecological role within the crust is still unknown. Other ecosystems where these fungi are found also have no external carbon or nitrogen sources, and these fungi are not known chemolithotrophs (rock degrading) (Gorbushina, 2007; Teixeira et al., 2017). Therefore, their sources of nutrients seem to be limited to their autotrophic microbial neighbors in these same ecosystems, which many have observed but never experimentally shown (Gorbushina et al., 2005; Gostinčar et al., 2012; Grube et al., 2013; Muggia et al., 2013). Due to what we know about the metabolic capabilities of these fungi, where we find them, and their evolutionary relatives the lichen-forming Verrucariales, they are more than likely using carbon and nitrogen fixers in their environment for nutrition acquisition.

### Endosymbiotic Bacteria of Fungi

While the concept of endosymbiosis has been around for many years, evidence of fungi containing endosymbiotic bacteria has been a relatively recent discovery (Bonfante, 2003; Kobayashi & Crouch, 2009). Most focus of bacterial-fungal endosymbiosis has been on the mycorrhizal fungi which associate with the roots of plants (Bianciotto et al., 2000; Lumini et al., 2007; Mondo et al., 2012). Even with more time and improved techniques, the diversity of fungi containing endosymbiotic bacteria has largely focused on plant-associated fungi (Bastías et al., 2020; Hoffman et al., 2013; Izumi et al., 2006; Pakvaz & Soltani, 2016; Ruiz-Herrera et al., 2015; Shaffer et al., 2017). Only very recently has a non-plant associated saprotrophic fungi, *Mortierella* spp., been identified to contain an endosymbiont (Büttner et al., 2021). Overall, the main role of bacteria in these symbioses has been in the direct production, or inducing the production, of a chemical to enhance survival of the fungus, with the implication that the bacteria is creating something for the benefit of the fungus and not necessarily benefiting the bacteria.

The scientific community’s attention on obtaining axenic cultures for fungal type strain deposition has probably reduced the number of identified endosymbiotic bacterial-fungal symbioses. Additionally, overuse of antibiotics to ensure clean cultures for genome sequencing likely also contributes to this issue. Many microbes are considered unculturable, but it remains possible that these microbes require the presence of their microbial neighbors for survival. The more we dedicate our time and resources to culturing non-laboratory or “unculturable” fungi, the more likely we will find fungi with uncurable endosymbionts.

### Methylobacterium’s metabolic capabilities and their roles in symbioses

*Methylobacterium* is a Gram negative alphaproteobacterial genus of the order Rhizobiales, now called Hyphomicrobiales. Also known as pink facultative methylotrophs (PPFMs), they are characterized by their ability to utilize one-carbon molecules such as methanol as a carbon source (Chistoserdova et al., 2003). They are prominent members of the plant phyllosphere (Kutschera, 2007), with certain taxa capable of root nodulation (Jourand et al., 2004; Sy et al., 2001); however, they are ubiquitously found in all ecosystems (Hiraishi et al., 1995). Metabolically, members of this genus have been known to perform nitrogen fixation (Jourand et al., 2004; Madhaiyan et al., 2015; Raja et al., 2006), aerobic anoxygenic photosynthesis (Atamna-Ismaeel et al., 2012; Csotonyi et al., 2010; Tang et al., 2018), and phytohormone production (Ivanova et al., 2001; Ivanova et al., 2000; Koenig et al., 2002). These metabolic capabilities make them ideal partners for broader interactions within a microbial community, as they have the capability to produce a nitrogen source, growth stimulating compounds, and perform the first step of photosynthesis creating ATP for themselves.

As their ubiquitous nature would imply, many *Methylobacterium* spp. have been identified across multi-species symbioses. Importantly, *Methylobacterium* have been identified as a part of newly recognized lichen holobionts (Cardinale et al., 2006; Cardinale et al., 2008; Erlacher et al., 2015; Hodkinson & Lutzoni, 2009; Jiang et al., 2020). Their role in the lichen symbiosis is currently unknown, but many still hypothesize that the unique metabolic capacity of *Methylobacterium* spp.’ allow them to engage in nitrogen fixation, some role possibly associated with their nodulation capabilities, and phytohormone production for growth-promoting capabilities (Erlacher et al., 2015; Grube et al., 2009; Hodkinson & Lutzoni, 2009). Of particular interest to this study, *Methylobacterium* spp. have also been found within the biological soil crust consortium (Csotonyi et al., 2010; Tang et al., 2018). The ecological importance of *Methylobacterium* spp. within the biological soil crust seems to be significant, as enriching for *Methylobacterium* spp. via activation of aerobic anoxygenic photosynthesis increases the growth of the biological soil crust (Tang et al., 2018). It is likely that *Methylobacterium* spp. could play a pivotal role in the biological soil crust consortium, with their wide assortment of metabolic capabilities at the ready for microbial interaction optimization.

We describe here a novel polyextremotolerant fungal species and its presumptive *Methylobacterium* spp. endosymbionts. Full characterization of the fungus has been performed in lieu of its genome, which is still being sequenced and analyzed. Additional experiments have been performed to understand the basis for this fungal-bacterial symbiosis, to determine if it is a true endosymbiotic interaction, and the ecological niche this fungal-bacterial symbiosis might occupy within the biological soil crust consortium.

## Methods

### General methods

#### Microbial strains and growth conditions

This chapter describes one novel fungal species and its two *Methylobacterium spp*. symbionts. The novel species of fungi described here is currently named JF2 08-2F Crusty and will from here on be referred to as Crusty. The closest alignment to this isolate’s (ITS) fingerprint region is *Neophaeococcomyces* sp. Currently, it does not have a designated species name, as its genome sequencing and annotation is still being processed at the Joint Genome Institute (JGI). Once its genome and transcriptome are fully sequenced, they will be deposited at the Department of Energy (DOE) JGI’s Mycocosm website under the designated species name, and at the National Center for Biotechnological Information (NCBI). The type strain will also be deposited to the Westerdijk institute for access to the greater scientific community.

Crusty is grown in malt extract medium (MEA; composition in **Table 1**) at room temperature in an Erlenmeyer flask at 1/10^th^ the volume of the flask, shaking at 200 rpm. *Methylobacterium* spp. symbionts of Crusty are called JF2 08-2LP Light Pinky and JF2 08-2DP Dark Pinky, and the combination of these two in culture is called JF2 08-2P Pinky (from here on called Light Pinky, Dark Pinky, and Pinky respectively). According to the results of nucleotide similarity search using the Basic Local Alignment Search Tool (BLASTn) against the 16S ribosomal RNA (rRNA) sequence database at NCBI, both Light Pinky and Dark Pinky were 100% identical with an E-value of 0.0 to multiple *Methylobacterium* species including: *Methylobacterium oryzae* strain 94-A4, *Methylobacterium phyllosphaerae* strain CBMB27, and *Methylobacterium radiotolerans* strain PPFM28. *Methylobacterium* spp. are maintained in MEA as well with the same conditions stated above.

**Table 1:** Media compositions of media used in this study.

| Media Name | Acronym | Composition (L<sup>-1</sup>) |
| --- | --- | --- |
| Malt Extract Agar | MEA | 20 g Dextrose<br>20 g Malt Extract<br>2 g Peptone<br>15 g Agar |
| Minimal | MN | 10 g Dextrose<br>50 mL 20x Nitrate salts<br>1 mL Hutner's Trace Elements |
| Minimal + Vitamins | MNV | 10 g Dextrose<br>50 mL 20x Nitrate salts<br>1 mL Hutner's Trace Elements<br>1 mL Vitamin Mix to MN |
| Yeast Extract Peptone Dextrose | YPD | 20 g Dextrose<br>20 g Peptone<br>10 g Yeast Extract<br>20 g Agar |
| Bold's basal medium | BBM | (Connon, 2007) |
| Additives | Volume/L | Composition |
| 20X Nitrate Salts/MN salts | 50 mL | 120 g NaNO<sub>3</sub> (remove for "MN salts")<br>10.4 g KCl<br>10.4 g MgSO<sub>4</sub>·7H<sub>2</sub>O<br>30.4 g KH<sub>2</sub>PO<sub>4</sub> |
| Hutner's Trace Elements | 1 mL | 2.2 g ZnSO<sub>4</sub>·7H<sub>2</sub>O<br>1.1 g H<sub>3</sub>BO<sub>3</sub><br>0.5 g MnCl<sub>2</sub>·4H<sub>2</sub>O<br>0.5 g FeSO<sub>4</sub>·7H<sub>2</sub>O<br>0.17 g CoCL<sub>2</sub>·6H<sub>2</sub>O<br>0.16 g CuSO<sub>4</sub>·5H<sub>2</sub>O<br>0.15 g Na<sub>2</sub>MoO<sub>4</sub>·2H<sub>2</sub>O<br>5 g EDTA (Disodium)<br>100 mL ddH<sub>2</sub>O |
| Vitamin Mix | 1 mL | 10 mg biotin<br>10 mg pyridoxin<br>10 mg thiamine<br>10 mg riboflavin<br>10 mg p-aminobenzoic acid (PABA)<br>10 mg nicotinic acid<br>100 mL ddH<sub>2</sub>O |

Additional organisms used in this study were *Saccharomyces cerevisiae* strain BY4741, algae *Chlorella sorokiniana* strain UTEX 1230, and an additional *Methylobacterium* sp. wild isolate named P3 that was isolated from the North Antelope Valley Parkway canal in Lincon, NE. *S. cerevisiae* is grown in Yeast Peptone Dextrose medium (YPD; composition in **Table 1**) with the same flask and shaking conditions as described above. *C. sorokiniana* is either grown up in a carbonless medium called Bold’s basal Medium (BBM), or TAP which contains carbon, and is grown under 12 hr light/dark cycles with shaking at 200 rpm.

#### Fungal isolation and identification methods

Fungi were isolated from a biological soil crust located in Jackman Flats British Columbia, Canada (52.931383 N, 119.372059 W). Soil samples were taken from the top 2 cm of the biological soil crust. A 0.1 g portion of the crust was re-suspended in 1 mL of water, ground with a sterile micropestle, and diluted with a dilution factor (DF) of 10 until they reached 10,000x dilution. Each dilution was then spread out onto two different MEA petri plates containing either no antibiotics or containing: Ampicillin (100 mg/L), Chloramphenicol (50 mg/L), Gentamycin (10 mg/L), and Cycloheximide (100 mg/L). The plates were then incubated in a Percival light incubator at 23 °C with a 12 hr light/dark cycle and examined daily using a dissection microscope to check for small black colonies. Once a potential black colony was seen, half of it was removed and transferred to a new MEA (no antibiotics) petri plate. Plates were examined daily, because even in the presence of antibiotics many unwanted fast-growing microbes would grow on the plates and cover up the slower growing polyextremotolerant fungal colonies. Once a pure culture of each isolate was grown up (approximately 2 weeks), they were preserved in 30% Glycerol and stored at -80 °C.

DNA sequencing of amplified internal transcribed spacer (ITS) sequences was used to identify the isolates. DNA was extracted using the Qiagen DNeasy Powersoil DNA extraction kit. Primers used to isolate the ITS region were: ITS1- (5’-TCC GTA GGT GAA CCT GCG G-3’) and ITS4- (5’-TCC TCC GCT TAT TGA TAT GC-3’) (White et al., 1990). A BioRad MJ Mini Personal Thermal Cycler was used, with the program set as: 1) 95 °C for 5 minutes; 2) 94 °C for 30 seconds; 3) 55 °C for 30 seconds; 4) 72 °C for 1 minute and 30 seconds; 5) Return to step 2, repeat 35 times; 6) 72 °C for 5 minutes; 7) END. Amplified products were then checked via gel electrophoresis in 1% agar run at 80 V for 1 hr. Isolated ITS regions were sequenced using the Eurofins sequencing facility. Subsequently, a nucleotide similarity search using BLASTn was performed against the NCBI database to identify their potential taxanomical matches.

#### Phylogenetic Analysis of Crusty’s ITS Sequence

To further refine the identity of newly isolated strains, we generated a multi-loci gene tree using ribosomal ITS1, ITS2, LSU, SSU and 5.8S genetic markers. Because these markers are commonly sequenced through amplicon or Sanger sequencing, there is a higher availability of closely related species with representative markers than closely related species with whole genome sequences available for comparison. Fifty-seven sequences of strains from the NCBI and UNITE databases were collected based on the BLAST results of Crusty (Abarenkov et al., 2010). The program ITSx *v.1.1b* (Bengtsson-Palme et al., 2013) was used with the save_all tag to pull out the ITS1, ITS2, LSU, SSU and 5.8S regions from each read. As the data on NCBI were all collected using a variety of different primer and PCR set-ups, this program allows us to capture as much of the ribosomal regions as possible. The program MUSCLE *v.5.1* (Edgar, 2004) was used to align each fasta file containing ribosomal gene region. Clipkit *v.1.3.0* (Steenwyk et al., 2020), another program to help with overall alignment and trimming strategy, was used to retain parsimony informative sites that are known to be more phylogenetically informative. A python script (combine_multiseq_aln.py found in PHYling_united utilities folder https://github.com/stajichlab/PHYling_unified) was used to combine all aligned sequences into a fasta sequence. This python script also takes the length of each region into consideration as it builds the larger fasta file from a separate text file. The fasta files aligned and trimmed by MUSCLE and clipkit were each used and run with the nucleotide tag. Phylogenetic analysis was performed using IQ-Tree *v.2.2.0* (Minh et al., 2020), which was used in two main ways. Initially 1000 bootstrap Ultrafast (-b 1000) was used while also calling IQ-Tree automatic selection of substitution models for each ITS and rDNA region, specified as a partition. Standard evolutionary rates for branch lengths were used for each partition. The details along with the length of each region is added into a partitioning NEXUS file for multi-gene alignments and used to run IQ-Tree a second time and generate a composite ML phylogenetic tree. To reduce the computational burden that occurs with including a large number of organism ribosomal sequences, we used the relaxed hierarchical clustering algorithm, automatic model selection and best-fit partitioning scheme (Lanfear et al., 2014) were performed using the option “-m MF+MERGE”.

#### Bacterial isolation and identification methods

Bacterial isolates were obtained after streaking out Crusty from frozen stocks. If antibiotics are not present in the media used to streak out a frozen stock of Crusty, then pink bacterial colonies always form. The difference in colony color allowed the isolation of the two bacteria. Identification of these bacteria was done by extracting DNA as explained above, and then using PCR to amplify their 16S ribosomal DNA (rDNA) region using the primers: 8F (5’-AGA GTT TGA TCC TGG CTC AG-3’), and 1492R (5’-GGT TAC CTT GTT ACG ACT T-3’) (Turner et al., 1999). A BioRad MJ Mini Personal Thermal Cycler was used for performing the PCR, with the program set to 1) 95 °C for 5 mins; 2) 95 °C for 1 min; 3) 51 °C for 1 min; 4) 72 °C for 2 mins; Go to step 2, repeat 30 times; 5) 72 °C for 10 mins; 6) END. PCR products were run on a 1x TAE 1% agarose gel electrophoresis at 80V for one hour for verification. Sequencing of the PCR products was performed by MWG Eurofins using the 8F primer. The obtained sequences were analyzed using BLASTn nucleotide similarity search against the NCBI nr/nt database, and the closest matching bacterial species were determined.

#### Phylogenetic Analysis of Methylobacterium spp. using 16S rDNA Sequences

Phylogenetic analysis of Light Pinky and Dark Pinky was performed using their 16S rDNA marker region. We downloaded all 16S rDNA sequences from the NCBI database using the *Methylobacterium* 16S rDNA Gene search. We also downloaded a selection of 16S rDNA sequences from *Microvirga* to use as an outgroup for the tree. Alignment of the 16S rDNA sequences was done in MEGAX using Muscle alignment’s standard parameters. After alignment, the ends of the 16S rDNA sequences were trimmed due to size differences across the various sequences. No trimming was performed within the sequences beyond the end regions. On the very rare occasion that there was a strain with an insert, these regions were kept because it would be important for the further analysis to differentiate the closely related strains. A maximum likelihood tree was reconstructed in MEGAX using the GTR+G+I model with 1000 bootstrap values.

#### Methylobacterium spp. Genome Assembly and Annotation

Since the clarity of bacterial identification using the 16S rDNA sequence was not enough to obtain exact species identification, we performed sequencing of the whole genome of the two *Methylobacterium* spp. we isolated from Crusty’s culture: Light Pinky and Dark Pinky. DNA extraction of the isolates was done in the same way as described above for Crusty. The resulting genomes were sequenced using the Microbial Genome Sequencing Center (https://www.migscenter.com) using their 300 Mbs sequencing service. Resulting sequences were uploaded to the Galaxy web platform for computational analyses (Afgan et al., 2018). Bacterial genomes were aligned on Galaxy using the standard functions of Unicycler (Wick et al., 2017). Then, automatic annotation was performed on Galaxy using Prokka (Seemann, 2014). BLAST databases for Light Pinky and Dark Pinky’s genomes were also created on Galaxy using blastdbn (Camacho et al., 2009; Cock et al., 2015). Individual proteins were identified on the Dark Pinky genome using BLASTp which was ran on Galaxy (Camacho et al., 2009).

#### Initial Microscopy and Plate Imaging

Initial microscopic images were taken using an Olympus BX51 on the 40x objective lens, with an iPhone 6s against the ocular lens. Cells of JF2 08-2F Crusty were grown up as described above for 10 days. Then 8 μL of the grown cells was aliquoted onto a microscope slide with a 1.5 mm coverslip overtop. Plate imaging of Crusty, Light Pinky, and Dark Pinky was done using the iPhone 6s. Additional microscopy was done using the EVOS-fl at 4x-40x magnification.

#### Analysis of Budding Patterns

The protocol for observing the budding patterns of JF2 08-2F Crusty was derived from methods in (Mitchison-Field et al., 2019). A 1:1:1 ratio by weight of Vaseline, Paraffin, and Lanolin (VALAP) was combined in a glass bottle and heated to 115 °C to melt completely and kept at room temperature for later use. Heated solid MEA was aliquoted into a 50mL tube for agar slab making. Isolates were grown in liquid MEA for 10 days prior to inoculation of slides. First, the VALAP was brought back up to 115 °C to melt completely for application. Then 5 μL of the 10-day old cells were diluted in 95 μL of liquid MEA. Agar slabs of MEA were made by microwaving the 50 mL tube of solid MEA until it melted, then pipetting 1 mL of the hot agar into a 1 cm x 2 cm mold formed out of cut strips of silicone and laid down in a sterile petri dish. This agar slab was allowed to solidify and was then cut in half to be 1 cm x 1 cm. Both the cover slip and the slide were wiped down with ethanol to ensure clean and sterile growth conditions for the cells. 8 μL of the diluted cells was pipetted onto the center of the sterile slide, then one square of the agar slab was carefully placed on top of the cells in media, 8 μL of MEA was pipetted onto the top of the agar slab, and the coverslip was placed over the agar slab. Using a small paintbrush, the melted VALAP was carefully painted onto the gap between the coverslip and the microscope slide to seal off the coverslip. Finally, a 23-gauge needle was used to poke holes in the solidified VALAP to allow for gas exchange. The slide was then placed face down onto the inverted microscope EVOS fl. Once an adequate number of cells was observed in frame, the cells were allowed to settle for 2 hours before imaging began. Images were then automatically taken every 30 mins for 96 hours. Videos of the budding pattern were created using Adobe Premiere Pro.

#### XTT Assay Optimization of XTT:Menadione for Live Cell Growth

Planktonically grown microbial organisms can be measured using optical density (OD) for their cell growth cell growth. However, for organisms such as *Candida albicans* and Crusty who grow almost exclusively in biofilms, this form of measurement is highly inaccurate and, as such, much past work has been done to develop an indirect measurement of live cell growth. One such assay involves the use of sodium 3,3’-[l[(phenylamino)carbonyl]-3,4-tetrazolium]-bis(4-methoxy-6-nitro) benzene sulfonic acid hydrate (XTT) and Menadione for indirect measurement of live cells performing the TCA cycle via the production of red colored formazans (Berridge et al., 2005). This assay works by measuring trans-plasma membrane electron transport via reduction of menadione by XTT (Berridge et al., 2005). Although it is an indirect measurement, it is meant to be used as a comparison of the same organisms in different conditions allowing for an accurate comparison when no other means are viable.

Concentrations of XTT and menadione were optimized to measure live growth of JF2 08-2F Crusty (Antachopoulos et al., 2006). XTT is suspended in 1x PBS and menadione is suspended in acetone. A stock solution of XTT was made to the concentration of 0.5 mg/mL. Solutions of menadione were made up to the concentrations of 300 mM, 200 mM, and 50 mM. These solutions were then combined in a 20:5 XTT:menadione ratio and tested against a serial dilution of Crusty which was serial diluted with a dilution factor of 1.2 to allow for a gradual gradient of nine dilutions from 0-4.3x diluted. These dilutions were made by starting with 600 μL of cells in the 0x tube, and 100 μL of media in each dilution 1.5 mL tube. Then 500 μL of the previous dilution was added to the next dilution, mixed, and continued to the next dilution. Finally, 400 μL of fresh media was added back to the tubes to allow for a final volume of 500 μL for a proper volume to XTT:menadione mixture ratio of 500:25 (Moss et al., 2008). After the addition of the XTT:menadione, the cells were allowed to incubate at room temperature for 18 hours. The tubes were then centrifuged at 10,000 g, and 100 μL of the supernatant was aliquoted into a 96-well plate and measured at 460nm.

Comparison of the results of the individual XTT:menadione ratios to the serial dilution was performed. Linear regression analysis was performed on these values, and the optimal combination has the highest R^2^ value.

### Phenotypic Characterization of the Fungus Crusty

#### Carbon Source Utilization of Crusty

Carbon utilization of each isolate was determined using a BioMerieux ID C32 carbon utilization strip (Cat. No. 32200-1004439110). These strips have 30 different carbon sources in individual wells, one well with no carbon source for a negative control, and one well with esculin ferric citrate (**Table 2**). The esculin (β-glucose-6,7-dihydroxycoumarin) ferric citrate assay was originally used to identify Enterobacteria (Edberg et al., 1977), but was co-opted to identify *Cryptococcus neoformans* via melanin production and esculin degredation (Edberg et al., 1980).

**Table 2:** Carbon Utilization of Crusty. (n=3, 3 biological replicates)

| *Carbon Source* |  | *# 1* | *# 2* | *# 3* | *Average* | *Score* |
| --- | --- | --- | --- | --- | --- | --- |
| *D-galactose* | GAL | 4 | 4 | 4 | 4 | + |
| *D-sorbitol* | SOR | 5 | 4 | 5 | 4.7 | + |
| *Actidione (cycloheximide)* | ACT | 3 | 4 | 4 | 3.7 | + |
| *D-xylose* | XYL | 3 | 3 | 4 | 3.3 | V |
| *D-saccharose (sucrose)* | SAC | 5 | 5 | 5 | 5 | + |
| *D-ribose* | RIB | 2 | 3 | 1 | 2 | V |
| *N-acetyl-glucosamine* | NAG | 5 | 5 | 5 | 5 | + |
| *Glycerol* | GLY | 4 | 4 | 5 | 4.3 | + |
| *Lactic acid* | LAT | 2 | 2 | 2 | 2 | V |
| *L-rhamnose* | RHA | 3 | 4 | 4 | 3.7 | + |
| *L-arabinose* | ARA | 2 | 2 | 1 | 1.7 | - |
| *palatinose* | PLE | 3 | 4 | 4 | 3.7 | + |
| *D-cellobiose* | CEL | 5 | 5 | 5 | 5 | + |
| *Erythritol* | ERY | 2 | 1 | 1 | 1.3 | - |
| *D-raffinose* | RAF | 5 | 5 | 5 | 5 | + |
| *D-melibiose* | MEL | 4 | 3 | 3 | 3.3 | V |
| *D-maltose* | MAL | 2 | 2 | 2 | 2 | V |
| *sodium glucuronate* | GRT | 3 | 3 | 3 | 3 | V |
| *D-trehalose* | TRE | 2 | 1 | 1 | 1.3 | - |
| *D-melezitose* | MLZ | 5 | 5 | 5 | 5 | + |
| *potassium 2-ketogluconate* | 2KG | 4 | 4 | 3 | 3.7 | + |
| *potassium gluconate* | GNT | 2 | 2 | 1 | 1.7 | - |
| *methyl-αD-glucopyranoside* | MDG | 2 | 1 | 1 | 1.3 | - |
| *levulinic acid (levulinate)* | LVT | 2 | 2 | 2 | 2 | V |
| *D-mannitol* | MAN | 4 | 4 | 5 | 4.3 | + |
| *D-glucose* | GLU | 5 | 5 | 5 | 5 | + |
| *D-lactose* | LAC | 1 | 1 | 1 | 1 | - |
| *L-sorbose* | SBE | 4 | 4 | 4 | 4 | + |
| *Inositol* | INO | 4 | 4 | 4 | 4 | + |
| *glucosamine* | GLN | 1 | 1 | 1 | 1 | - |
| *no substrate* | No | 1 | 1 | 1 | 1 | - |
| *Esculin ferric citrate* | ESC | + | + | + | + | + |

The inoculated strips were kept in a plastic box container with a lid and lined with moist paper towels to reduce drying. The initial inoculum of cells was prepared in 25 mL of MEA shaking at room temp for 5 days. Inoculation of the strips was done according to the instructions provided by the vendor, and each strain was inoculated in three separate strips for triplicate replication. Cells were diluted to the kit requirement of McFarland standard #3 (McFarland, 1907) before starting the inoculum. Growth in the ID strip lasted 10 days before evaluation. Growth in each well was observed and evaluated by eye.

Each well was compared to the negative control (no carbon) and the positive control (dextrose/glucose). Growth was evaluated on a scale of 1-5: 1 being no growth or resembling the growth of the No carbon negative control, and 5 being the highest growth resembling the growth on glucose. Three replicates were performed, and their averages were taken to determine the kit’s final scale of: +, V, and -. For this scale: 1-1.9 was “-”, 2.0-3.6 was “V”, and 3.7-5.0 was “+”.

#### Nitrogen Source Utilization of Crusty

Nitrogen utilization tests were performed using ten different nitrogen conditions. 100 mM was the concentration used for all compounds that contained one nitrogen atom per molecule: Proline, Ammonium tartrate dibasic, Serine, Sodium Nitrate, Glycine, Glutamate, and Aspartate; 50 mM was the concentration used for Urea because it has two atoms of nitrogen per molecule; 1% w/v of Peptone was used as a positive control; and no nitrogen was added as a negative control. This concentration of nitrogen sources was used because Crusty is typically grown on MEA or YPD which contain 0.2% and 2% peptone respectively. Total nitrogen content of Peptone is around ∼15%, with our standard YPD media using 2% peptone that equates to 0.3% total nitrogen in the media which is 150-200 mM of nitrogen source (peptone’s nitrogen content varies by batch). We decreased that molarity to 100 mM for ease of solution making but is meant to represent the amount of nitrogen source between MEA and YPD which is their standard growth media. Liquid minimal media (MN) with MN salts (not 20x Nitrate salts) (**Table 1**) was used with the varying nitrogen sources to ensure that no alternative nitrogen source would be available to the fungi. Fungi were first grown up in liquid MEA for 5 days at room temperature to reach maximum density. One mL of cells was removed and washed three times with water. 195 μL of each nitrogen source-containing medium was added to six wells in a 96-well plate, for six replicates, and 5 μL of the washed cells was added to each well. 200 μL of each medium was also added to one well each without cells to blank each condition, because the different nitrogen sources created different colors of medium. After 7 days of incubation, growth of Crusty on each nitrogen source was measured via the XTT assay because extensive biofilms form on the bottom of the wells which are difficult to disrupt physically. Specifically, it was assumed that there was 200 μL total volume in each well and the XTT assay was scaled to that volume, where 10 μL of the XTT:Menadione was added to each well and the assay was carried out as described above.

#### UV-resistance of Crusty

To assess UV resistance, we used the UVP HL-2000 HybriLinker UV crosslinker as our source of UV light, which has a UV wavelength of 254 nm. Lower wavelengths (100-280 nm) are of the UV-C range, are considered ionizing radiation and are the most detrimental to living organisms, but are completely blocked by the ozone layer (Molina & Molina, 1986; Schreier et al., 2015). Therefore, using this wavelength we are able to push our organisms beyond the UV limits found in their natural habitat and test extreme amounts of UV exposure.

The fungi were inoculated in 25 mL of MEA in a 250 mL Erlenmeyer flask and let grow under shaking conditions at 200 rpm for 10 days at room temperature to reach maximum density. This culture was then washed, and serial diluted to 0x, 10x, and 100x dilution in water to ensure that as many individual cells were being exposed to the UV light as possible. 100 µL of each dilution was then spread out onto 6 MEA plates, using a glass spreader. Three plates were kept as the control growth, to compare to the three other plates which were exposed to the UV light. Experimental plates were placed inside of the crosslinker with their lids taken off. Then the plates were exposed to 120 seconds of UV light from a distance of 9.5 cm to the light source at 10,000 μJ/cm^2^ (254 nm) (Frases et al., 2007). We then wrapped all plates in aluminum foil and placed them in the Percival light incubator set at 23 °C for 2 days. Placing UV-exposed cells in complete dark after exposure is essential for preventing light-dependent photoreactivation of induced lesions (Weber, 2005). After 2 days the plates were removed from the aluminum foil and left in the incubator for 5 more days before final observations. To determine whether a particular isolate was resistant to UV exposure, the growth of the isolate exposed to UV was compared to the control growth.

#### Metal-Tolerance of Crusty

Metal resistance is a relatively universal trait in many polyextremotolerant fungal species. Due to the under-studied nature of this particular characteristic in fungi from biological soil crusts, we decided to test if any of our isolates were resistant to any heavy metals which would indicate possible bioremediation capacity. In order to test metal resistance, we used the antibiotic disc method by aliquoting metal solutions onto paper discs and observing zones of clearance. Metals and concentrations used are listed in **Table 3**. For testing, 5 µL of each metal solution was aliquoted onto a dry autoclaved Whatman filter paper disc which was created using a standard hole puncher. These discs were allowed to air dry and kept at 4 °C for up to a week. Initial growth of the fungal isolates was done in 25 mL of MEA, shaking at 200 rpm for 5 days at room temperature. We spread 100 μL of each fungal isolate onto 100 mm sized MEA plates using a glass spreader to create a lawn. Using flame sterilized forceps, our metal paper discs were placed onto the center of the petri dish on top of the fungal lawn and lightly pressed down to ensure the metal disc was touching the plate completely. These plates were placed in the Percival light incubator at 23 °C with a 12 hr light/dark cycle for up to 2 weeks. Once a zone of clearing was clearly visible amongst the fungal growth (1-2 weeks), the zone of clearing was measured in cm. Generally, large zones of clearing indicated sensitivity to the metal, whereas zones of reduced size were consistent with resistance to the metal.

**Table 3:** Average zone of clearing for all metals tested on Crusty. (n=3, 3 biological replicates)

| Metal | Zone of clearing (cm) |
| --- | --- |
| 0.5 M $\text{CoCl}_2$ | 3.0 |
| 470 mM $\text{AgNO}_3$ | 2.8 |
| 1 M $\text{FeSO}_4$ | 0.0 |
| 2 M $\text{LiCl}_2$ | 0.0 |
| 1.5 M $\text{CuCl}_2$ | 1.4 |
| 1.5 M $\text{NiCl}_2$ | 4.7 |
| 10 mM $\text{CdSO}_4$ | 4.5 |

#### Temperature Growth Range of Crusty

To determine the temperature resistance range and optimal growth temperature for JF2 08-2F Crusty, we grew the fungus at 4 °C, 15 °C, 23 °C (i.e., ambient room temperature), 28 °C, 37 °C, and 42 °C. JF2 08-2F Crusty was first grown up in 25 mL of MEA for 10 days at room temperature to maximum density. One mL of cells was removed, and a 10x serial dilution was made from 0x to 100,000x, using pre-filled 1.5mL tubes with 900 µL of MEA and adding 100 µL of the previous tubes each time. Five µL of each serial dilution was spotted onto a square MEA plate which allowed us to determine the carrying capacity of each isolate at the different temperatures. Plates were kept at their respective temperatures for 7 days before observations were made, however the 37 °C and 42 °C incubators required cups of water inside of them to prevent the plates from dehydrating. Plates grown in 42 °C and 37 °C were allowed to grow at room temp for up to a week to determine if the isolates died at these temperatures or if their growth was just arrested.

### Experiments to decipher the relationship between Crusty and its Methylobacterium spp. symbionts

#### Determination of Endosymbiosis via 16S rDNA Amplification Post-Chloramine-T Treatment

Multiple methods obtained from various other fungal-bacterial endosymbiont manuscripts were tested on Crusty and its *Methylobacterium* spp. co-occurring bacteria. One such experiment was to surface-sterilize the fungal cells using a combination of Tween-20, hydrogen peroxide, and Chloramine-T as described by (Mondo et al., 2012). Exact methods from Mondo et al. were followed; however, to make up for the clumped nature of Crusty, manual disruption of the clumps was done using a micropestle against 800 μL of Crusty that was grown up as previously stated above. This would ensure that as many cell surfaces were exposed as possible, to reduce the likelihood of bacterial cells being trapped between cell clumps. The treatment was then used on the “post-pestled” cells, and 16S PCR amplification as stated above was performed on pre- and post-Chloramine-T treated cells to determine if surface sterilization removed all signs of bacterial presence indicating non-endosymbiosis, or if a 16S signature remained, most likely indicating endosymbiosis.

#### Confocal Microscopy to Observe Bacterial Presence Inside Fungal Cells

To observe the presence of the bacterial symbionts inside of Crusty, we used a combination of Propidium Iodide (PI), SYTO9, and Calcofluor staining. PI and SYTO9 are considered a live/dead stain for bacteria where SYTO9 stains live cells in green (480nm/500nm) and PI stains dead cells in red (610nm). The protocol used for this method was done as follows, modified from (Partida-Martinez & Hertweck, 2005). Both PI and SYTO9 solutions were made up to 1.67 mM in DMSO, equal parts of each were added to a light-blocking 1.5 mL tube. 3 μL of that PI+SYTO9 mixture was added to 1 mL of washed (with water three times) Crusty cells that had grown up as described above but only for 6 days. The stains stayed in the tube of cells for 15 minutes before 2 μL of 1 mg/mL calcofluor was added to the cells right before visualization.

#### Antibiotics to Treat Crusty of its Methylobacterium Co-occurring Bacteria

In hopes of creating a clean strain of Crusty which did not contain any *Methylobacterium* spp. in its culture as a control, we attempted to use multiple antibiotics to remove the bacteria without killing the fungus. Antibiotics used were the β-lactam Ampicillin 100 mg/mL stock, peptidyl transferase Chloramphenicol 30 mg/mL stock, aminoglycoside Gentamycin 10 mg/mL stock, aminoglycoside Kanamycin 50 mg/mL stock, rRNA binding Tetracycline 100 mg/mL stock, and the fungicide Cycloheximide 100 mg/mL stock. To determine which antibiotic would be most effective against the *Methylobacterium* spp., we used the antibiotic disc method on the mixed culture of Pinky. Pinky was grown in MEA liquid for 5 days at room temperature, then 100 μL of the culture was spread out onto 18 MEA plates. Then filter paper discs were impregnated with 5 μL of one of the above listed antibiotics and were placed in the middle of the Pinky lawns. The plates were allowed to grow up for 8 days at 23 °C, then the zone of clearing caused by the antibiotic discs was observed. Once the antibiotic with the most defined zone of clearing was identified, we went on to use it on Crusty to try to clear out the *Methylobacterium* spp. from Crusty.

#### Gentamycin Treatment of Crusty, Pinky, and S. cerevisiae

Gentamycin ended up being the second most effective antibiotic treatment against the Pinky in our antibiotic disc trials. We decided to use Gentamycin as our first antibiotic to attempt to remove the *Methylobacterium* spp. from Crusty because it is bactericidal. Crusty, Pinky, and *S. cerevisiae* were grown up as described above, then 100 μL of the fungal cultures or 10 μL of the bacterial culture were added to 1.9 mL of MEA in a 24-well plate. In triplicate, each organism either had 20 μL of 0.3 mg/mL Gentamycin added, or nothing for a control. Then the cultures were allowed to grow at 23 °C for 7 days, after which the XTT assay as described above was used to determine the viability of cells after treatment.

#### Testing Fungal and Bacterial Ability to Grow in the Presence of Tetracycline

To attempt to obtain a Pinky-free Crusty and determine a dosage effect of the antibiotic Tetracycline on Crusty’s ability to grow, we exposed the fungus to increasing amounts of Tetracycline in liquid culture and then spread out the resulting growth on to plates with and without Tetracycline added. Crusty was grown up as described above, then 50 μL of cells were inoculated into size 25 tubes containing 5 mL of MEA. Each tube then received either 1 μL/mL, 2 μL/mL, 3 μL/mL, or 4 μL/mL of 100 mg/mL Tetracycline, 4 μL/mL of 100% Ethanol as a carrier control, or nothing as a standard control. This was done in triplicate tubes, and then they were put into a roller drum for 7 days. After 14 days of growth, 100 μL of each tube was spread out onto either a MEA plate with 1 μL/mL of Tetracycline added or no Tetracycline added. These plates were then allowed to grow up for 14 days, and then they were observed and imaged. To determine the effect of the Tetracycline treatment on Crusty, we then used ImageJ to calculate the pixel area of Crusty on each plate.

This same protocol was repeated using *S. cerevisiae* BY4741 and Light Pinky alone and in co-culture. This experiment allowed us to determine whether Tetracycline was affecting not only the bacteria, but also the fungus. We only used the highest concentration of Tetracycline, 4 μL/mL, for optimum effect and compared it to the Ethanol carrier control.

### Microbial Interaction Experiments

#### Tri-culture Experiments in Carbonless BBM

In lichens, the essential nutrient exchange process that is known is carried out by the photosynthetic partner providing a source of carbon to the fungal partner. We wanted to see if we could mimic this symbiotic phenomenon using Crusty, the algae *C. sorokiniana* UTEX 1230, and the addition of external Pinky. *C. sorokiniana* was chosen as the algal portion of this experiment because it is in the same class Tebouxiophyaceae as lichen forming algae of the genus *Trebouxia*. It was also chosen because of availability, it was the only algal culture in our collection at the time, and since it was related to *Trebouxioid* algae we believed it would be a comparable substitute for a *Trebouxia* spp.

This experiment was performed using the carbonless media Bold’s basal medium (BBM) (Connon, 2007) with and without its nitrogen source, which is sodium nitrate. All three organisms were grown up in their standard manner as described above, the cells were washed with BBM-nitrate three times, then 100 μL of each organism was inoculated into non-tissue culture treated tissue culture flasks either in monoculture, co-culture, or tri-culture. We used non-tissue culture treated tissue culture flasks because they have not been treated with vacuum-gas plasm, keeping their surface hydrophobic, which is optimal for fungal biofilm formation (Silva-Dias et al., 2015). The cells were then grown at 23 °C in a 12 hr light/dark cycle for 30 days. After 30 days, pictures of the entire flasks were taken and the XTT assay as described above was performed to determine the amount of active metabolism of Crusty in the various conditions. The XTT assay was not effective at determining the active metabolism of *C. sorokiniana*, but the images of flasks show where growth was capable or not capable.

#### Crusty-Pinky +/- Carbon, Nitrogen, Light

After performing the Tri-culture experiments there was uncertainty about how Crusty was capable of active metabolism when exposed to 30 days without a carbon source in the conditions that did not contain *C. sorokiniana*. Therefore, we decided to grow Crusty-Pinky (which is emphasized here as Crusty-Pinky due to Pinky’s importance) in BBM with and without 27.4 mM of mannitol, with and without sodium nitrate, and with and without the addition of 12 hours of light per day. This experiment was done to determine if light was affecting Crusty-Pinky’s ability to perform active metabolism in carbonless, nitrogenless medium. It was initially done to determine if the *Methylobacterium* spp. co-occurring with Crusty were capable of nitrogen or carbon fixation and therefore providing nutrients to Crusty allowing for active metabolism. However, this experiment was performed before we received the whole genomes of Light Pinky and Dark Pinky, where we discovered that Dark Pinky was capable of making bacteriochlorophyll and therefore capable of performing aerobic anoxygenic photosynthesis.

Crusty was grown up as described above, then an aliquot of cells was washed three times with water and let to sit in water for three hours. Then 20 mL of BBM either with or without 27.4 mM of mannitol and/or sodium nitrate was added to non-tissue culture treated flasks (25 cm^2^, 70 mL volume Falcon brand), where 100 μL of the washed cells were then added (n=3, 3 biological replicates for each condition). Flasks were placed inside of a Percival Scientific light incubator Model no.: CU-36L4, set to 23 °C with a 12-hour light/dark cycle. Flasks that were kept in the dark were kept in the same incubator but were wrapped entirely in aluminum foil to block all light. Cells were allowed to grow in these conditions for 14 days, and afterwards the XTT assay as described above was performed to determine active metabolism of Crusty-Pinky. Additionally, 100 μL of the culture was removed at day 7 and 14 to spread out onto MEA plates to determine if Pinky was growing or “escaping” Crusty, and if Crusty and/or Pinky was growing under each condition. Images of the bottom of the flasks were also taken at days 7 and 14 using an EVOS-fl, which has an inverted objective allowing us to not disturb the flasks for imaging.

#### Overlaying Algae over Methylobacterium spp. Spots

*Methylobacterium* spp. are known to enhance the growth of plants via the production of phytohormones (Ivanova et al., 2001). As such, we decided to determine if Light Pinky, Dark Pinky, their combination Pinky, and the North Antelope valley Parkway canal isolate P3, were capable of stimulating the growth of microscopic plants, aka algae. *Chlorella sorokiniana* has previously been identified as both responding to and producing its own auxins, which is why this strain was used in this experiment (Khasin et al., 2018). We tested this by spotting the bacteria onto MEA plates, then creating a semi-solid BBM overlay containing varying concentrations of *C. sorokiniana*.

The four bacterial cultures were first grown up as described above, then washed in fresh MEA three times. 5 μL of this culture was then spotted onto MEA plates nine times and repeated in triplicate plates for all three cultures with the intention to add algal overlays of five concentrations for each bacterial strain (45 plates total). The MEA plates with spotted bacteria were allowed to grow at room temperature for three days before the algal overlays were added. For the overlays, *C. sorokiniana* UTEX 1230 was grown up as described above, then washed three times with fresh BBM. This culture was then diluted with a 10x dilution from 0x-10,000x dilution in 10 mL aliquots. A semi-solid BBM mixture was made with 0.5% agar, and 9 mL of that mixture was added to 15 mL falcon tubes. 1 mL of the individual *C. sorokiniana* serial dilutions was added to the 9 mL of 0.5% agar BBM when the agar was approximately 40 °C. The BBM and *C. sorokiniana* mixture was then carefully poured over the bacterial spotted plates, as to minimize disturbances to the bacterial colonies. These plates were then grown at 23 °C with a 12-hour light/dark cycle for five days, after which they were observed and photographed.

## Results

### Description of Crusty

JF2 08-2F Crusty was isolated from Jackman Flats Provincial Park in B.C. Canada. Initial ITS sequencing was performed to obtain potential taxonomic matches. The highest BLAST match was 92.91% identity against “*Exophiala placitae* strain CBS 121716” (MH863143.1). Whole genome sequencing is still being performed. However, phylogenetic analyses using ITS revealed that JF2 08-2F Crusty is a novel species closely related to the genera *Phaeococcomyces* and *Neophaeococcomyces* (**Figure 1**). Morphological characterization of Crusty demonstrated that it has a very sand-like colony texture, allowing for the colonies to easily fall off a sterile loop when transferring to a plate, hence the name Crusty (**Figure 2A**). The colonies are relatively tall in the z-axis but hyphae do not tend to invade the agar itself. Crusty has a very dark black colony pigmentation, which is common to all polyextremotolerant fungi due to high melanin content in their cell walls (**Figure 2**). It also forms extensive biofilms on the air-liquid surface interface of flasks it’s grown in (**Figure 2C**).

**Figure 1:**
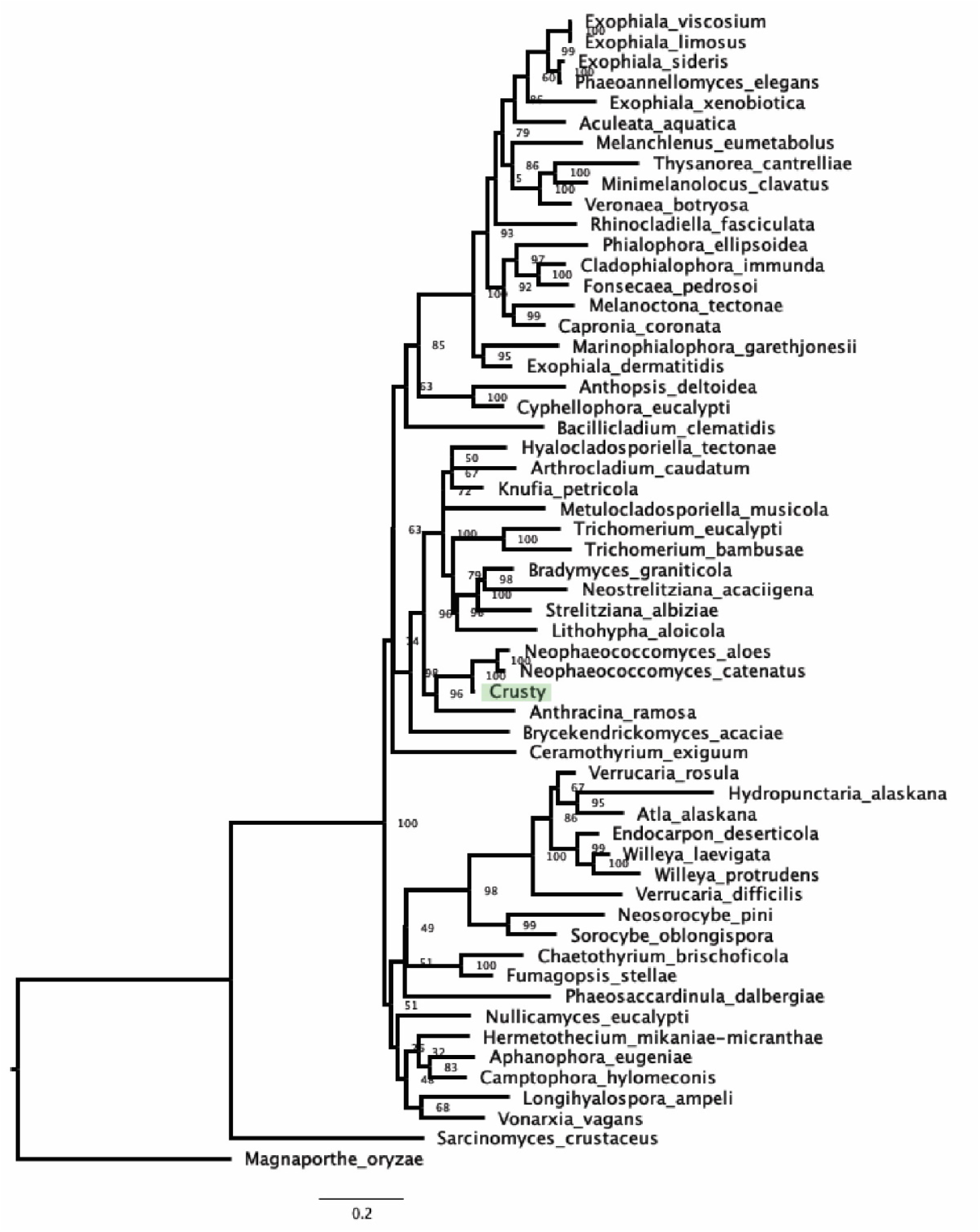
Maximum likelihood phylogenetic tree of Chaetothyriales using ITS sequences and 1000 bootstraps. The phylogenetic location of Crusty (green) within Chaetothyriales can be seen as closely related to *Neophaeococcomyces*, with a high confidence of a 96% bootstrap value. *Magnaporthe oryzae* was used at the outgroup.

**Figure 2:**
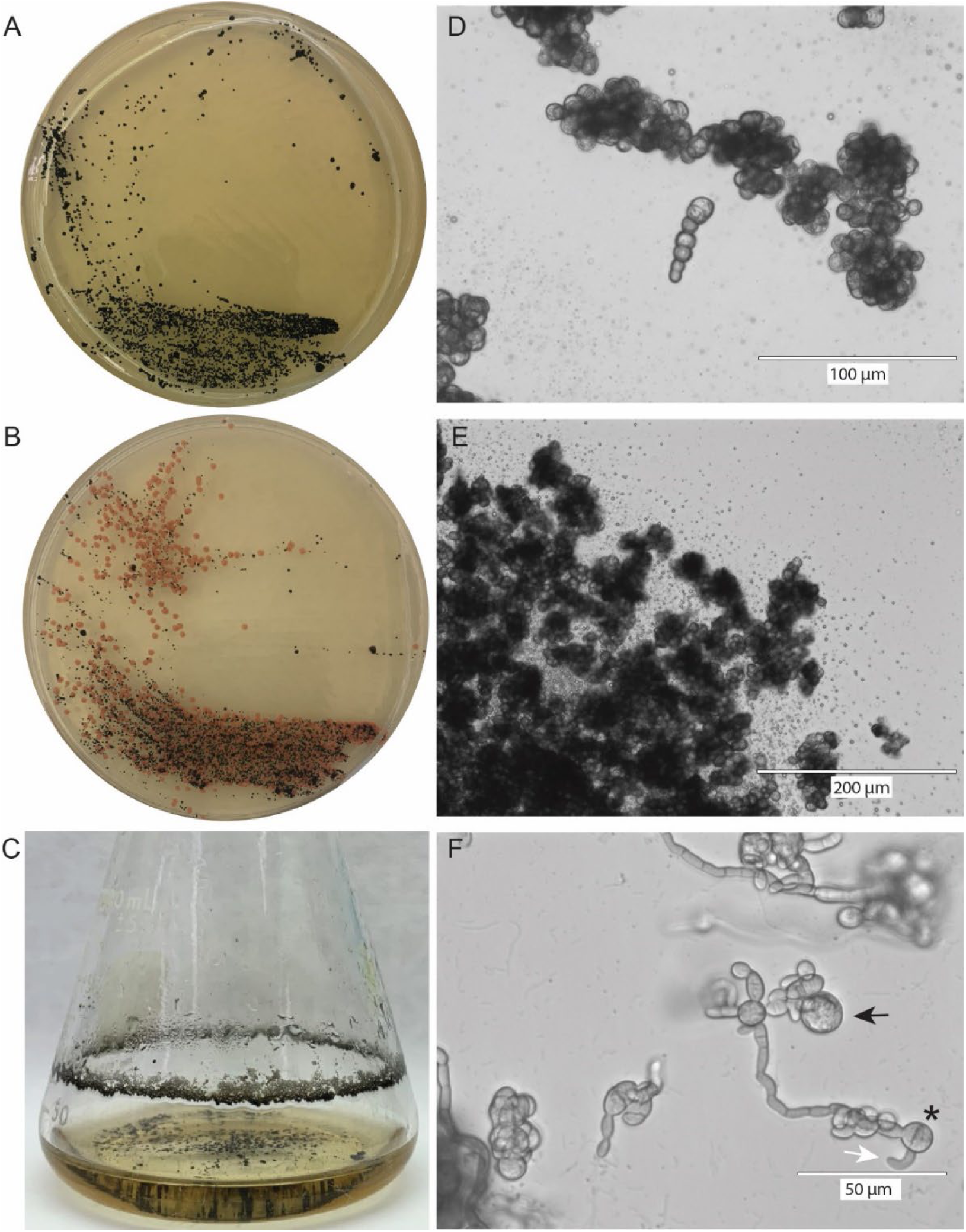
A) Colony morphology of Crusty grown on MEA + 10 mg/mL Tetracycline. Colonies are black, small, and no bacterial growth is present. B) Colony morphology of Crusty and Pinky when Crusty is grown from freezer stocks on MEA without antibiotics. Pinky colonies are able to grow as a result of the lack of antibiotics, due to burst cells from frozen stock. C) Liquid growth of Crusty in MEA. Crusty forms extensive biofilms at the air-liquid interface, that which forms distinct honeycomb-like patterns. In the liquid itself, Crusty does not grow planktonically but stays in its clumped morphology. D & E) Crusty cell morphology when grown on a plate of MEA. Perfectly round cells form, and high amounts of melanin are observed. E) Lipid droplets come out of large clumps of Crusty when the coverslip is placed over the cells. F) Crusty grown in BBM for 20 days showed unique morphologies: black arrows are large round cells; asterisks are multi-septated meristematic cells; white arrows are curved cells.

### The Many Faces (Cell Morphologies) of Crusty

Cells of Crusty are a polymorphic in nature, ranging from perfect spheres (**Figure 2D**) typical of the microcolonial fungi group of polyextremotolerant fungi, all the way to the formation of yeast and pseudohyphal cells (**Figures 2F & 3**). Other times, very large spherical cells can be seen with no observable septa, which are possibly chlamydospores (**Figure 3**; black arrows), and additional curved cells are frequently observed (**Figure 3**; white arrows). Crusty’s cell morphology also differs when grown in liquid medium versus growing on solid agar plates. In liquid it forms more pseudohyphae, and odd morphologies such as curved cells, meristematic cells, and large spherical cells are more commonly observed. Whereas, on solid media Crusty grows almost exclusively as perfect spherical cells in clumps and chains similar to *Knufia petricola*, and these cells are observed to release large amounts of lipid droplets from their surface (**Figures 2D & 2E**). Curved cells in fungi are usually a sign of chemotrophism attributed to the fungal cell’s attraction to a chemical compound. At this time, it remains unclear whether Crusty exhibits true chemotrophism and the nature of any possible chemical attractant is unknown. Additionally, Crusty’s cell division in liquid culture is similar to that of *Hortaea werneckii* and *Phaeotheca salicorniae*, in that it seems to both perform budding and binary fission in some aspects (**Figure 3B**), and also form meristematic cells and hyphae in other parts of budding cells (Mitchison-Field et al., 2019; Sterflinger, 2006) (**Figures 2F & 3;** asterisks).

**Figure 3:**
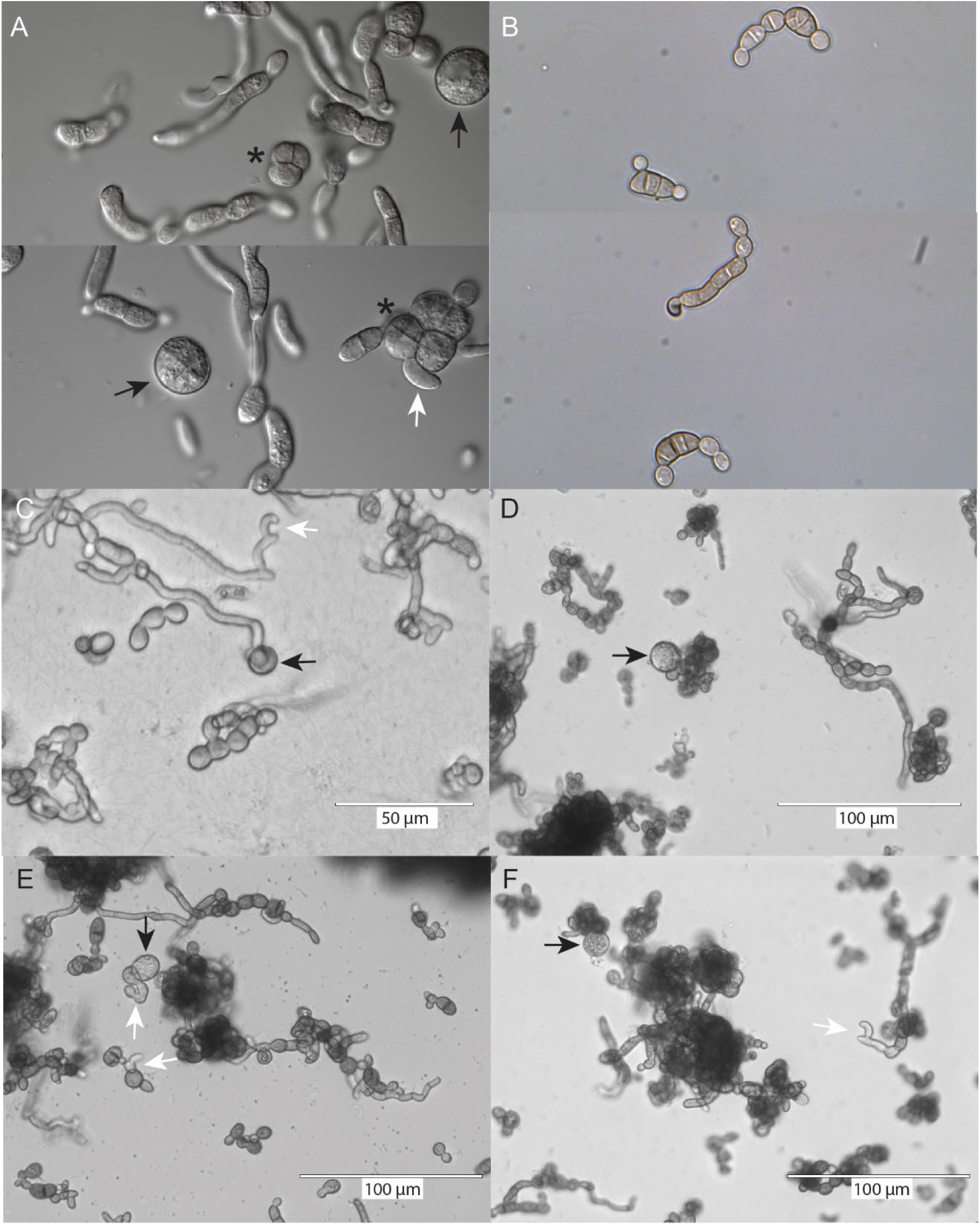
Various cell morphologies of Crusty cells. A & B) Crusty grown up in liquid MEA for 6 days. B) C) Crusty was grown up in BBM plus mannitol with a 12 hour light/dark cycle for 6 days. D, E, & F) Crusty was grown in BBM with no nitrate and added mannitol with a 12 hour light/dark cycle for D & F) 10 days and E) 17 days. White arrows point to curved cells; black arrow indicates large circular cells, possibly chalmydospores; asterisks indicate possible meristematic cell morphologies.

### Discovery of Methylobacteria Associated with Crusty

Typically, when growing out Crusty, it is not obvious that there are bacteria present because no bacterial colonies form (**Figure 2A**). If the cell wall has been disrupted in some way (i.e., resurrection of frozen stocks, bead beating), then colonies of both Crusty and Pinky will form on the plate when plating out cells (**Figure 2B**). This is why it was not initially apparent that this fungus even had bacterial symbionts. There are other strains in our collection from the same location that are inundated with *Methylobacterium* colonies when they are plated out, to the point where it is almost impossible to keep the fungus alive without the use of antibiotics to kill off the bacteria. Further evidence for the presence of bacteria associated with Crusty arose from the analysis of genomic DNA prepared for sequencing by the Joint Genome Institute (JGI). The extracted DNA had seemingly come from a sample that was a pure culture by eye, but results from quality controls revealed that there were two *Methylobacterium* spp. contaminating our sample. After performing our own 16S rDNA PCR again on what presumably was a pure culture of Crusty, we were able to replicate these results implicating that Crusty had co-occurring bacterial members that were not obvious when streaked on a petri plate.

### Spontaneous Albino (Pink) Mutant of Crusty

While Crusty was being domesticated in the lab, observations were made of pink colonies forming at random intervals that showed the same colony morphology and cell morphology as wild type Crusty (**Figure 4**). This is not a novel phenomenon for fungi of the *Knufia*, *Phaeococcomyces*, and *Neophaeococcomyces* clade as it was first observed by Katja Sterflinger’s group in *K. petricola,* with no follow-up however on the mechanism of its occurrence (Tesei et al., 2017).

**Figure 4:**
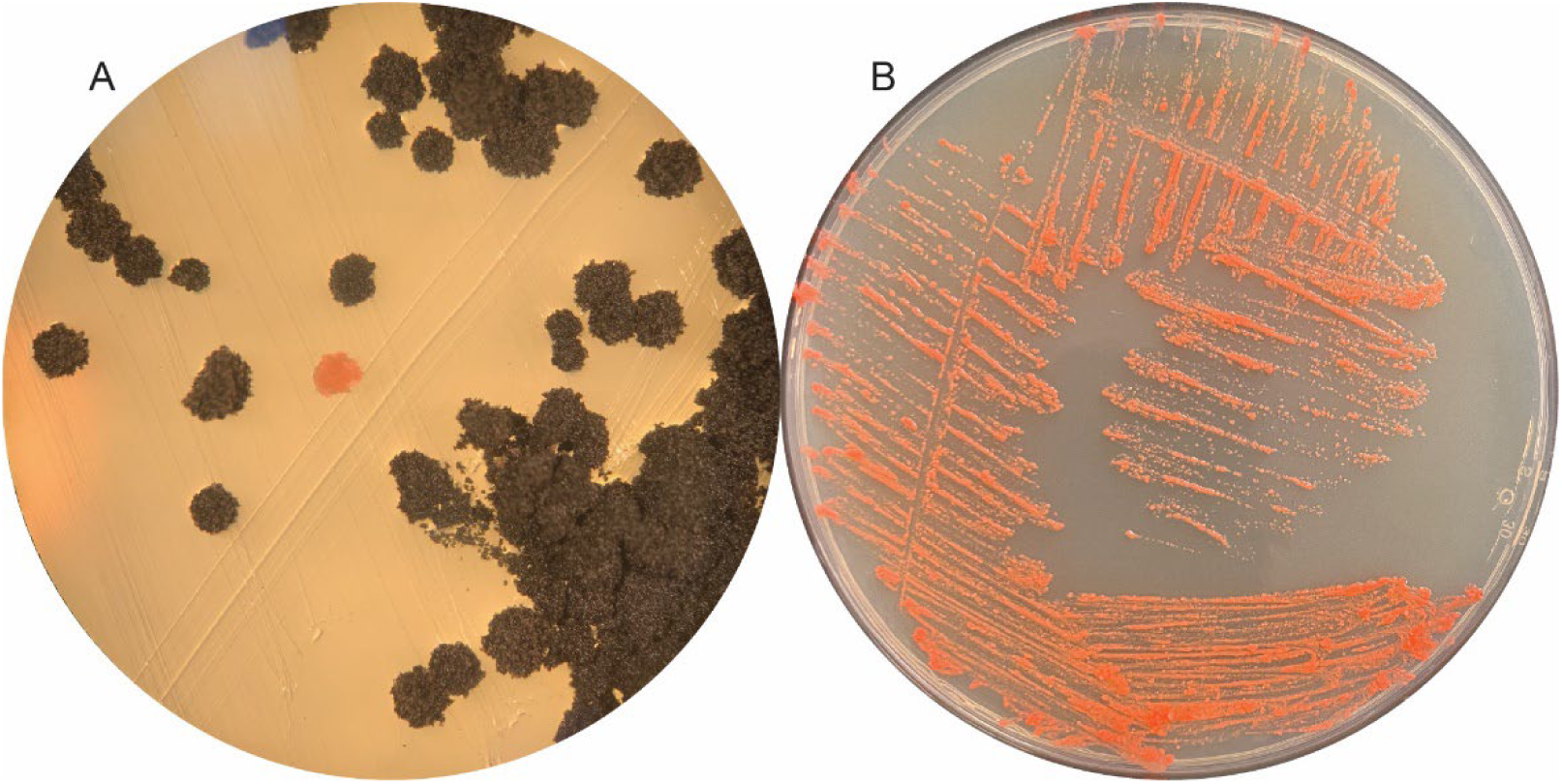
A) First instance of a pink albino colony of Crusty forming spontaneously on an MEA plate. B) Pure culture of the albino pink Crusty grown on MEA.

### Descriptions of Light Pinky and Dark Pinky, and Potential Metabolic Functions

Both of the bacteria found in association with Crusty are of the genus *Methylobacterium* according to their 16S rDNA and whole genome sequences. However, the species delineation between *Methylobacterium* spp. based solely on 16S rDNA is difficult to parse out (**Figure 5**). Therefore, assigning either JF2 08-2 LP Light Pinky or JF2 08-2 DP Dark Pinky to any known *Methylobacterium* spp. would require detailed genomic analyses. Nevertheless, we can still confirm that they are *Methylobacterium* spp., and with their genomes sequenced we will be able to understand what they are metabolically capable of as members of that genus. The relative abundance of these two bacterial strains within Crusty is unknown. It is even sometimes difficult to determine if they are both present simultaneously, if they are actually one strain, or if there is only one strain present using 16S rDNA PCR alone. Additionally, their genomes are highly similar, yet their morphologies remain distinctly different (**Figure 6**).

**Figure 5:**
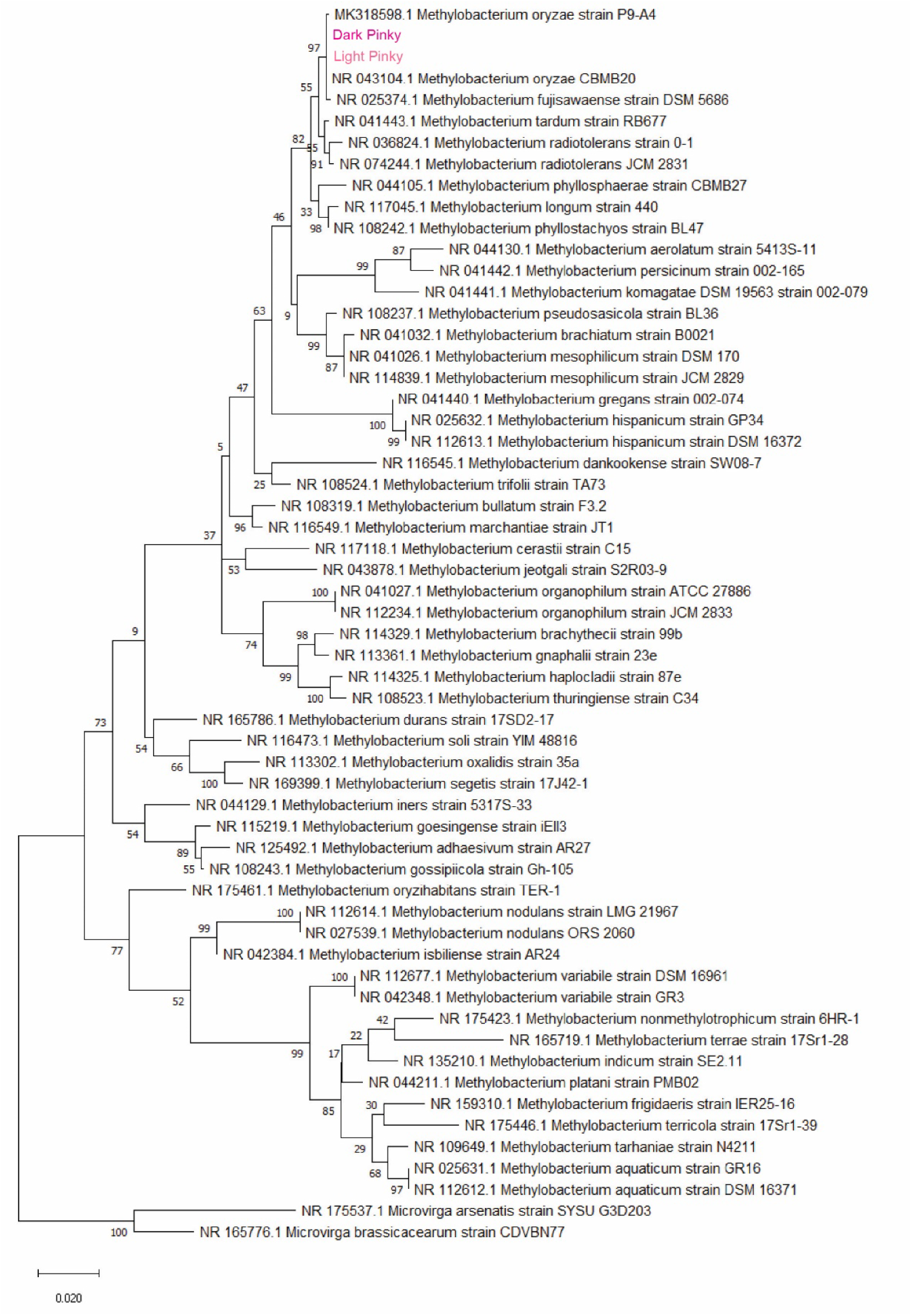
Maximum likelihood phylogenetic tree of the 16S rDNA region of *Methylobacterium* spp. with 1000 bootstraps. Values at nodes represent bootstrap values (%) of the nodes. Phylogenetic analyses confirm that Light Pinky and Dark Pinky are *M. oryzae*, but also closely related to *M. fujisawaense,* confounding the species name. *Microvirgra* species were used as the outgroup.

**Figure 6:**
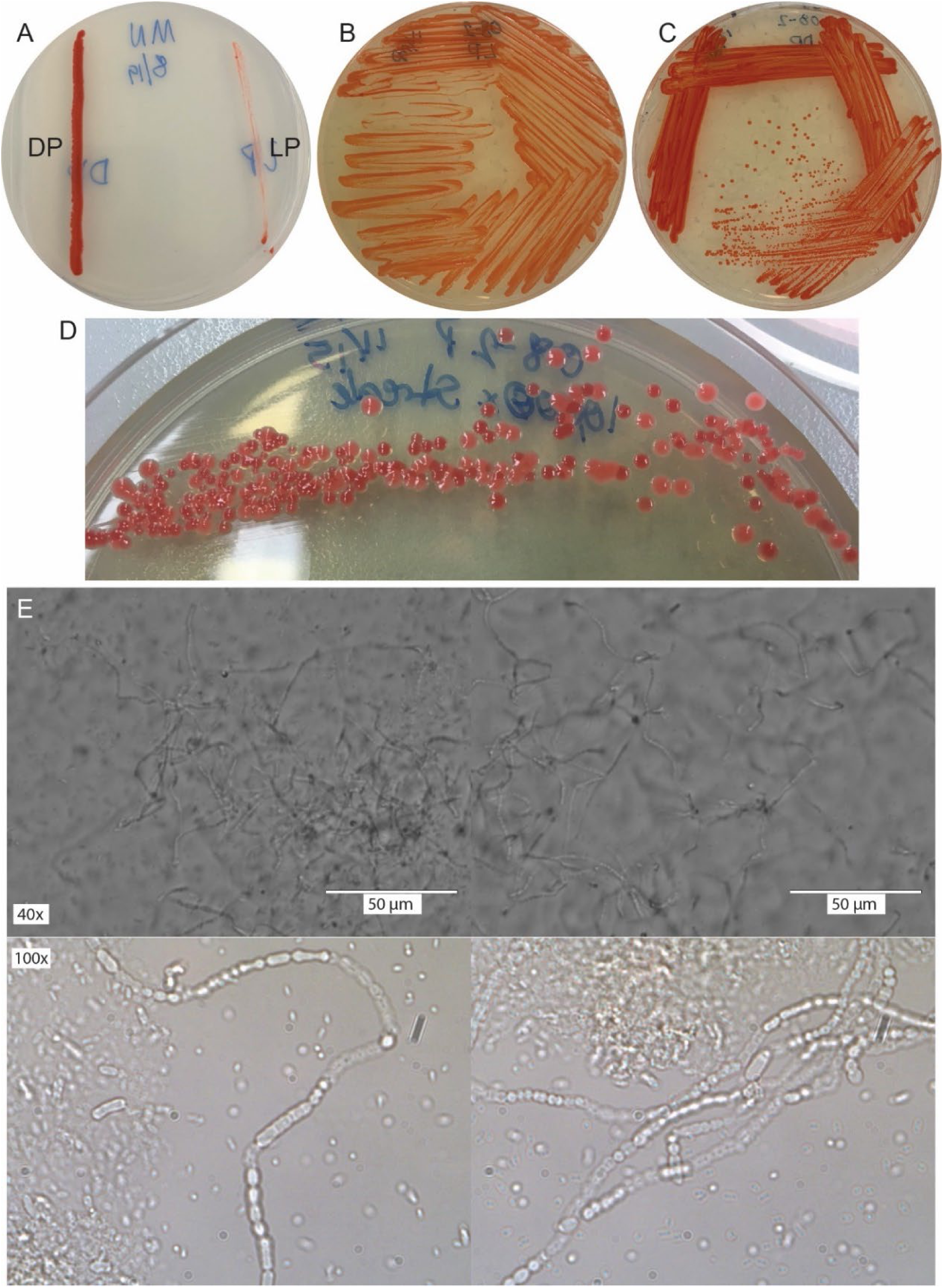
A) Dark Pinky (DP) and Light Pinky (LP) plated out on MN media together. On MN media it is obvious their color distinction, and the reduced growth by Light Pinky. B) Light Pinky grown on MEA. C) Dark Pinky grown on MEA. D) Colony morphologies of Light Pinky and Dark Pinky when growing out a 1000x diluted stock of their combined form Pinky, also shows their distinct color differences. E) Pinky control grown in BBM for 22 days, filamentation occurs.

Both Light Pinky and Dark Pinky are single-celled rod-shaped bacteria. In MEA, they do not readily form any filamentation or clumpy forms, although filamentation has been observed when grown in BBM (Data not shown). They are both pink in color due to the production of carotenoids, but have a distinct difference in their pigmentation, hence their difference in names. Light Pinky is lighter in color than Dark Pinky, where it is most obvious when they are grown on Minimal (MN) media (**Figure 6A**).

Regarding their metabolic capacity, genomic annotation suggests that neither strain is capable of nitrogen fixation as observed in Rhizobiales, for which there are three separate genetic pathways. However, in annotating these genes, we confirmed that they do contain ancestrally related genes involved in light-independent (dark) protochlorophyllide reductase (DPOR) used for anoxygenic bacterial photosynthesis (Fujita & Bauer, 2000). We were able to identify homologs of every bacteriochlorophyll (*bch*) gene required for bacterial photosynthesis from the Dark Pinky annotation (**Table 4**). Other genes essential for oxygen sensing that are used for both nitrogen fixation and anaerobic anoxygenic bacterial photosynthesis, such as the *fix* genes and other nitrogen fixation (*nif*) genes besides Nitrogenase iron protein 1 (*nifH*), were also found in the genome. These genes could be used to assist the bacteria in sensing an oxygen-less environment, which is essential for anaerobic anoxygenic photosynthesis, but not aerobic anoxygenic photosynthesis (Madigan, 1995).

**Table 4:**
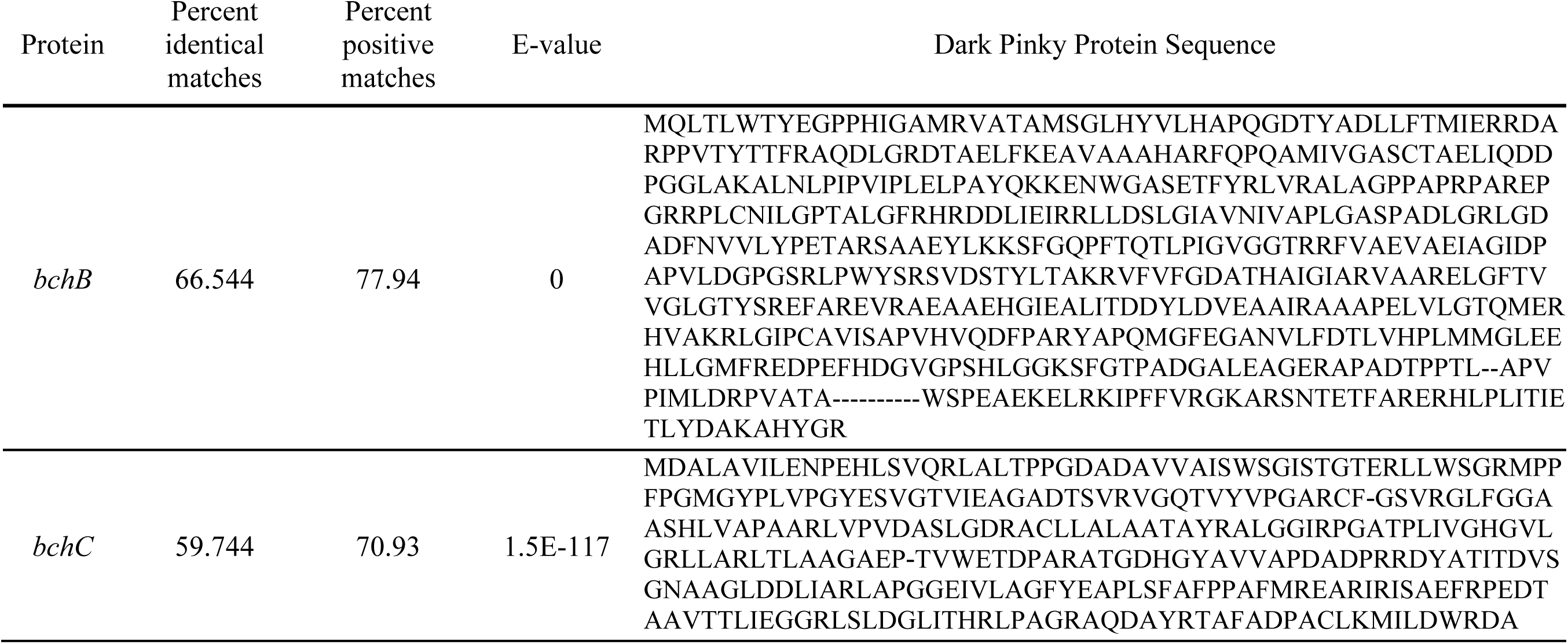
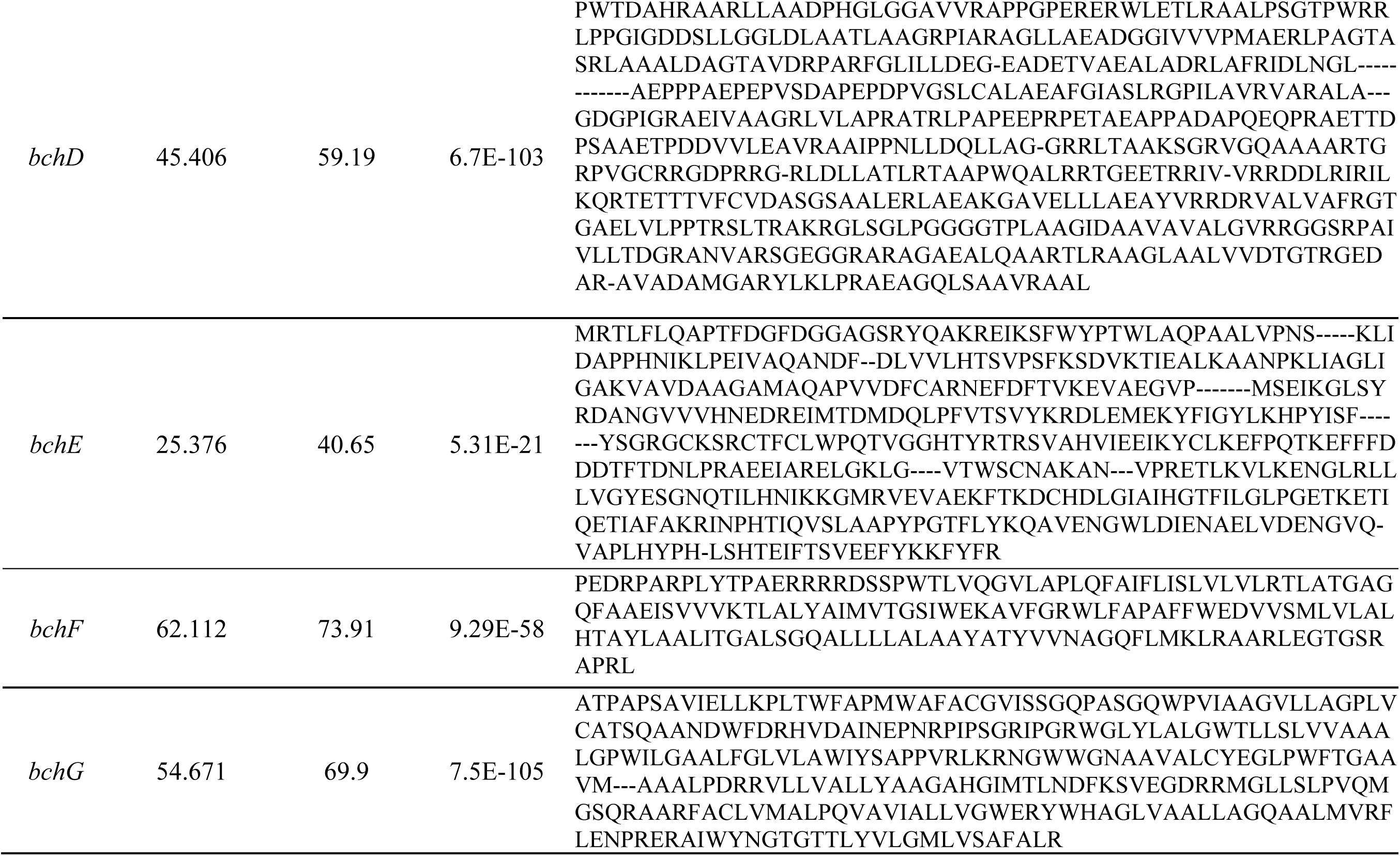
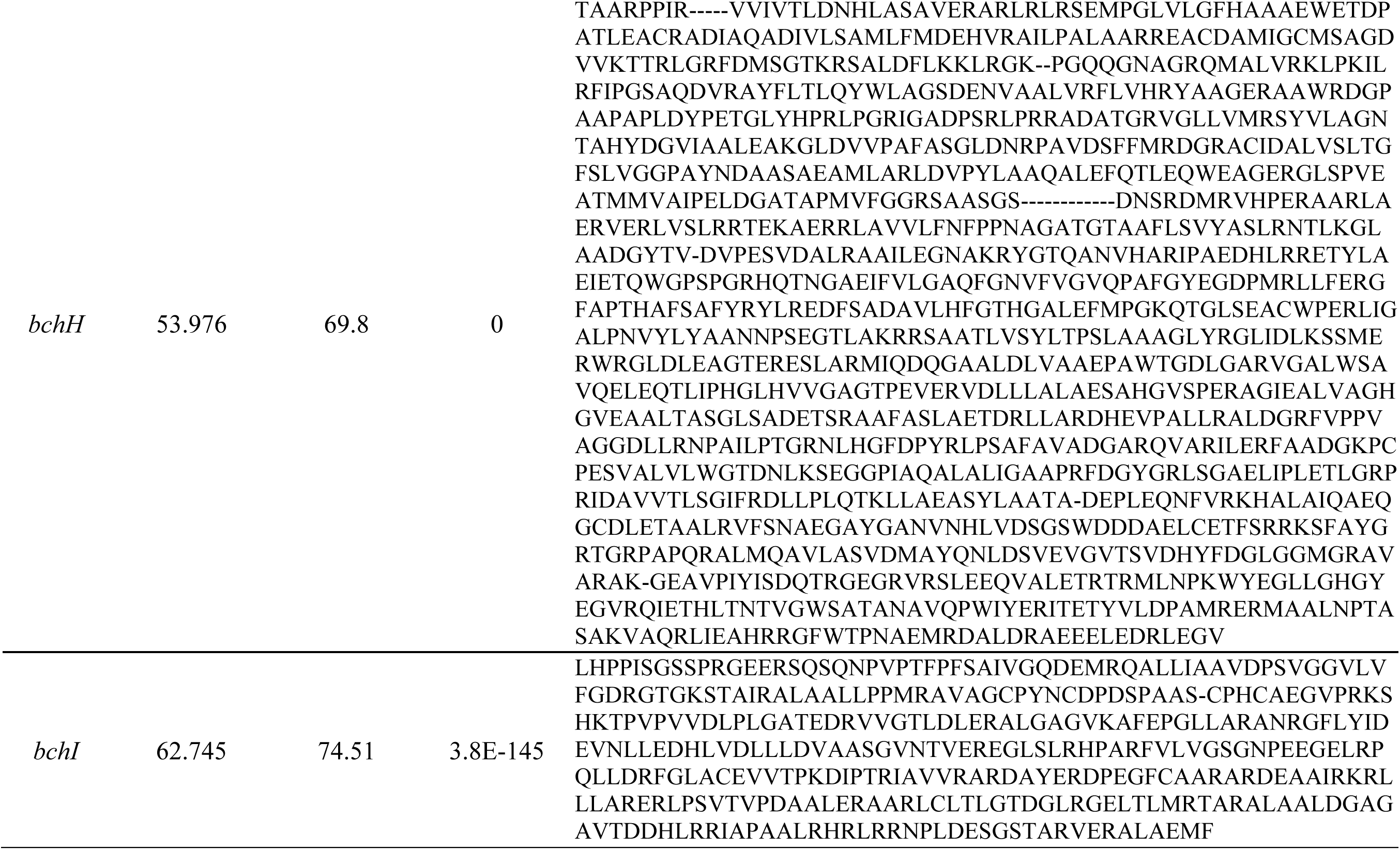
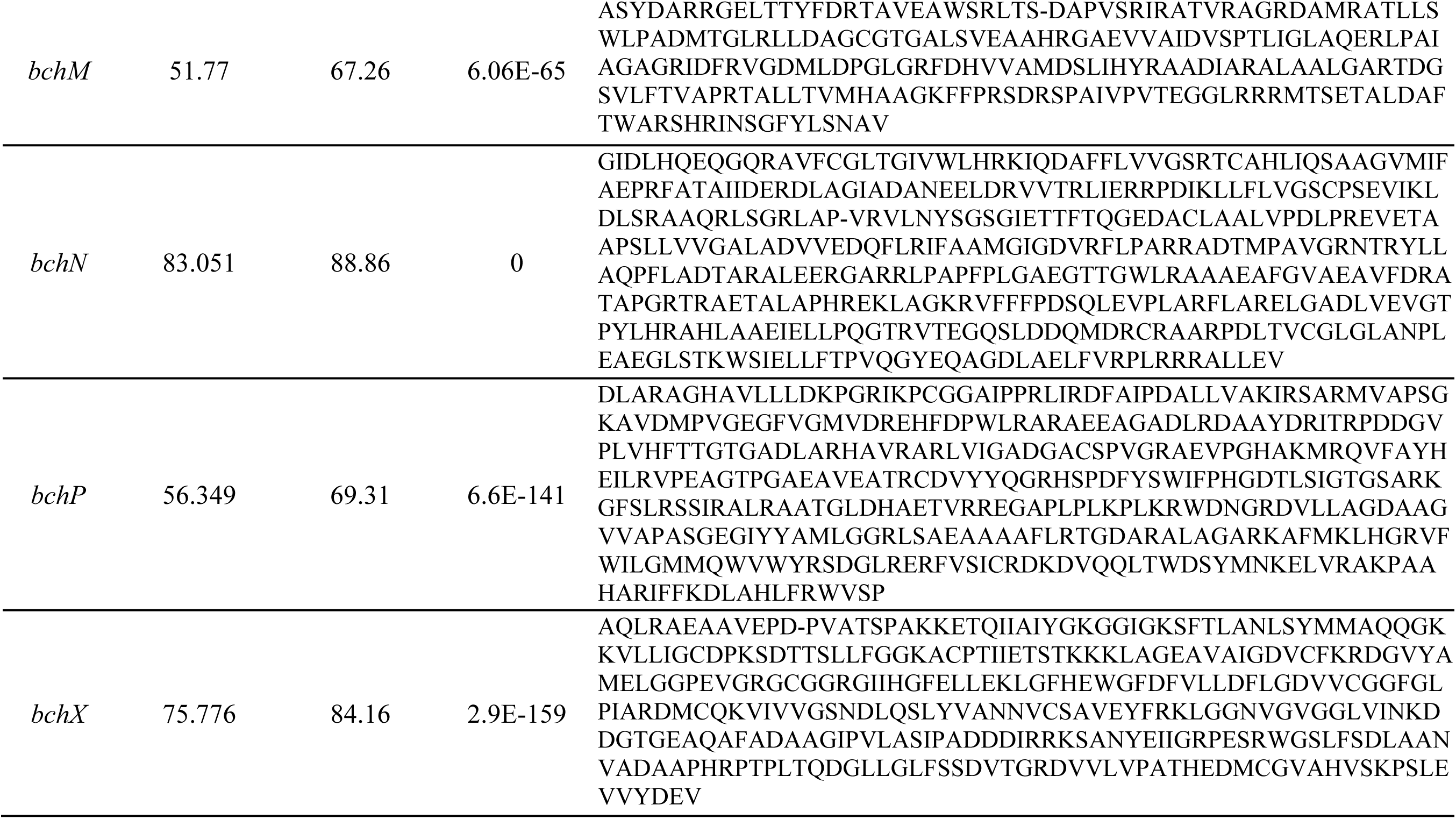
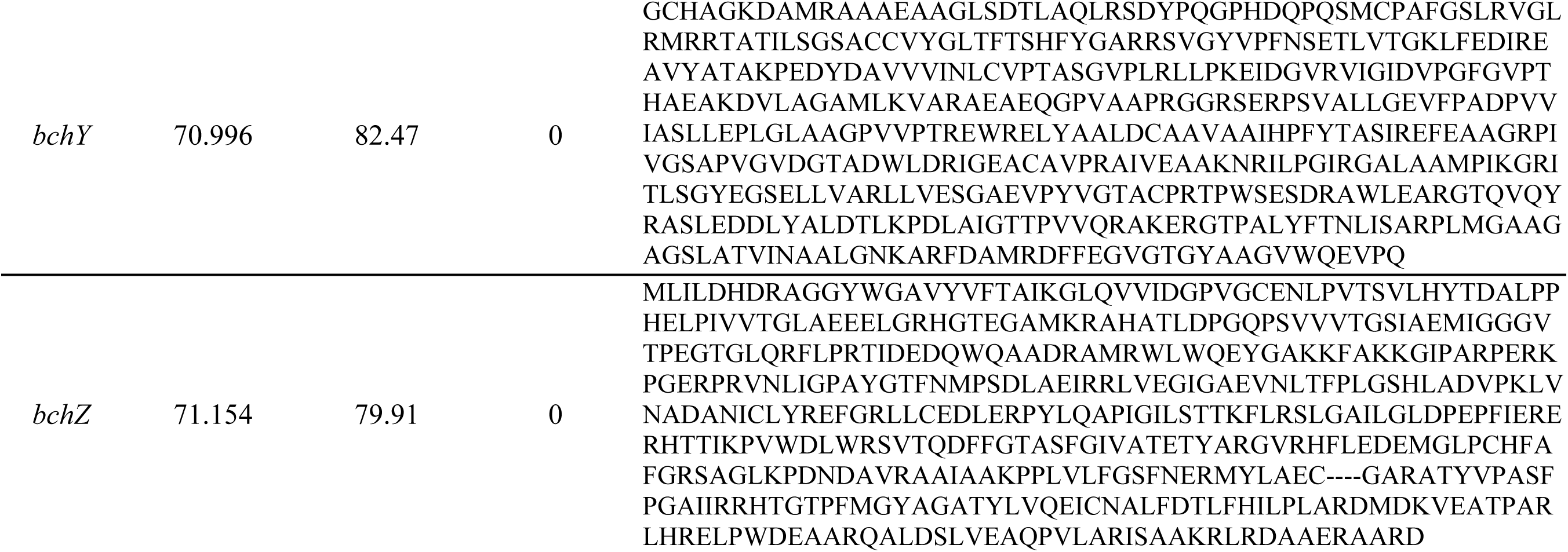
BLASTp results of all bch genes against the Dark Pinky genome. All proteins sequences were originally obtained from *Rhodobacter capsulatus,* except *bchN* came from *Methylobacter extorquens* and *bchB* came from *Cereibacter sphaeroides*.

### XTT Assay Optimization of XTT:Menadione for Live Cell Growth

Due to Crusty’s clumpy morphology and biofilm formation, performing any experiments that require a plate reader (OD reads) is challenging. However, many studies of other biofilm-forming or clumping organisms have implemented an alternative colorimetric assay, which utilizes live cells’ ability to transport electrons across their membrane during active metabolism as an alternative to direct OD reads (Berridge et al., 2005). Therefore, we decided to use this method to observe the growth patterns of Crusty in the variety of experiments we would impose upon it. This method needed to first be optimized for concentrations of the two substrates to Crusty before experiments could be performed.

Serial dilutions of Crusty were tested against the three different concentrations of XTT:Menadione of 0.5 mg/mL:50 μL, 0.5 mg/mL:100 μL, and 0.5 mg/mL:300 μL. Based on the resulting R^2^ values, the optimal XTT:Menadione concentration that was tested was 0.5 mg/mL:50 μL, and therefore was used for future studies (**Figure 7**).

**Figure 7:**
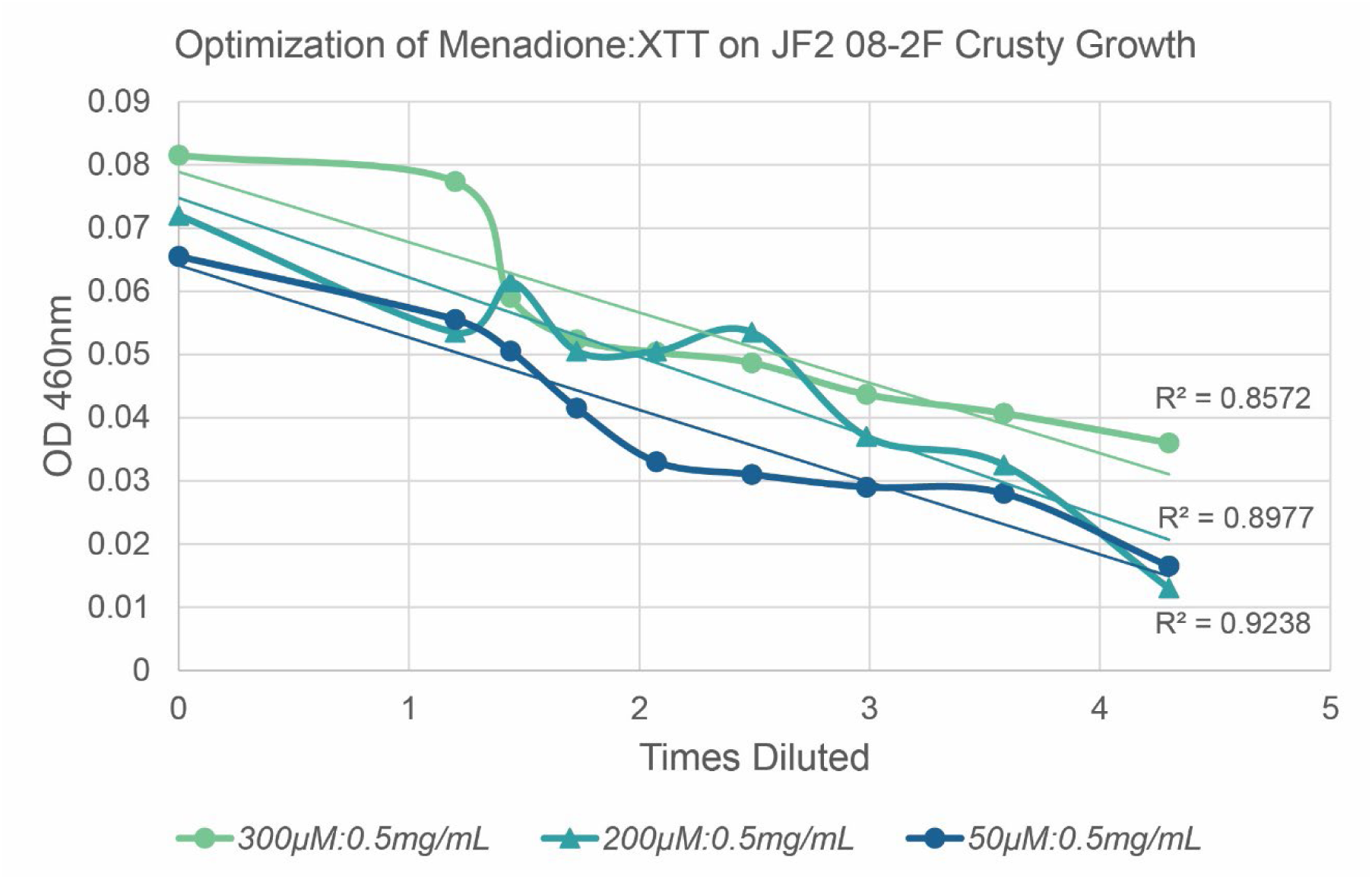
Optimization of the XTT assay used for multiple experiments. Specific amounts of Menadione to XTT were analyzed to determine which combination would most accurately represent the cell amount. The combination that had the highest R^2^ value would be the most accurate, which is shown here to be the 50 μM:0.5 mg/mL (menadione:XTT) combination at 0.9238.

### Phenotypic characterization of Crusty

#### Carbon Utilization of Crusty

Crusty was capable of utilizing 16 of the 30 carbon sources it was tested on in a positive manner, and 7 in a variable manner (**Table 2** & **Figure 8**). Overall, Crusty’s carbon utilization is similar to that of *E. viscosium* and *E. limosus* (Carr et al., 2022). However, two carbon sources that Crusty is capable of using that *E. viscosium* and *E. limosus* cannot use are D-saccharose and D-raffinose, and carbon sources that Crusty cannot use that the others can are L-arabinose, erythritol, D-trehalose, and potassium 2-ketoglutarate (**Figure 8**). Interestingly, we again see the presence of a difference in coloration with the growth of Crusty on N-acetyl glucosamine, making a reddish pigment. We were also able to confirm Crusty’s ability to produce DHN melanin with the esculin ferric citrate assay presenting a dark coloration (**Figure 8**)

**Figure 8:**
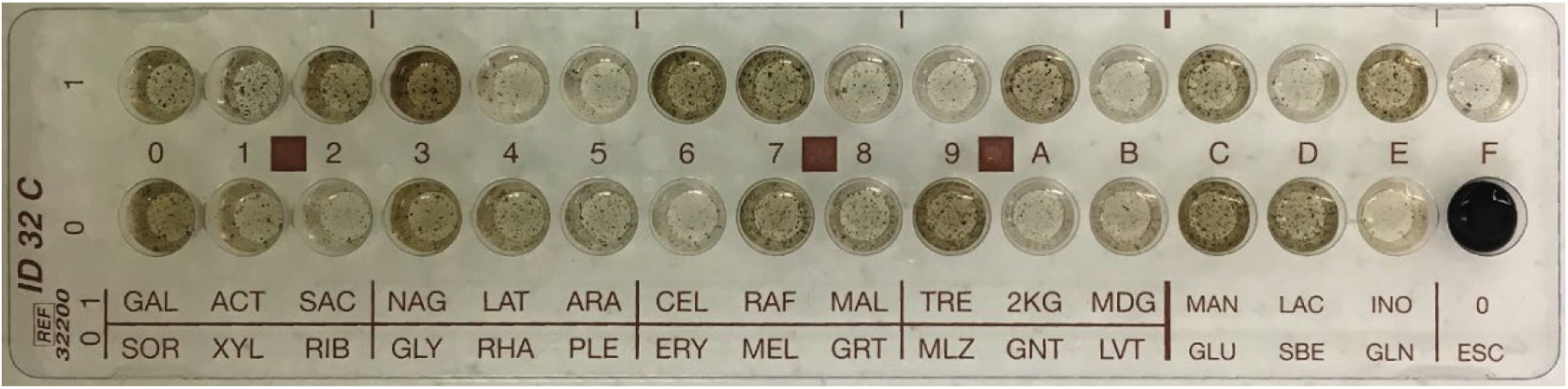
C32 strip results for Crusty. Darker wells indicate more growth which indicates more preference, except for ESC which is a chemical reaction not growth.

#### Nitrogen Utilization of Crusty

To investigate nitrogen source utilization of Crusty, we used MN media (**Table 1**) and dropped out the nitrate to add in our various nitrogen sources for testing Crusty’s ability to grow on them. According to the XTT assay and visual confirmation of growth, peptone remains Crusty’s preferred source of nitrogen, with urea and serine as close second preferable sources (**Figure 9**). Proline, glutamate, and aspartate were all similar in usage and the next preferred nitrogen source, followed by nitrate. Interestingly, the absence of a source seemed to promote more growth than did either glycine or ammonia (**Figure 9**).

**Figure 9:**
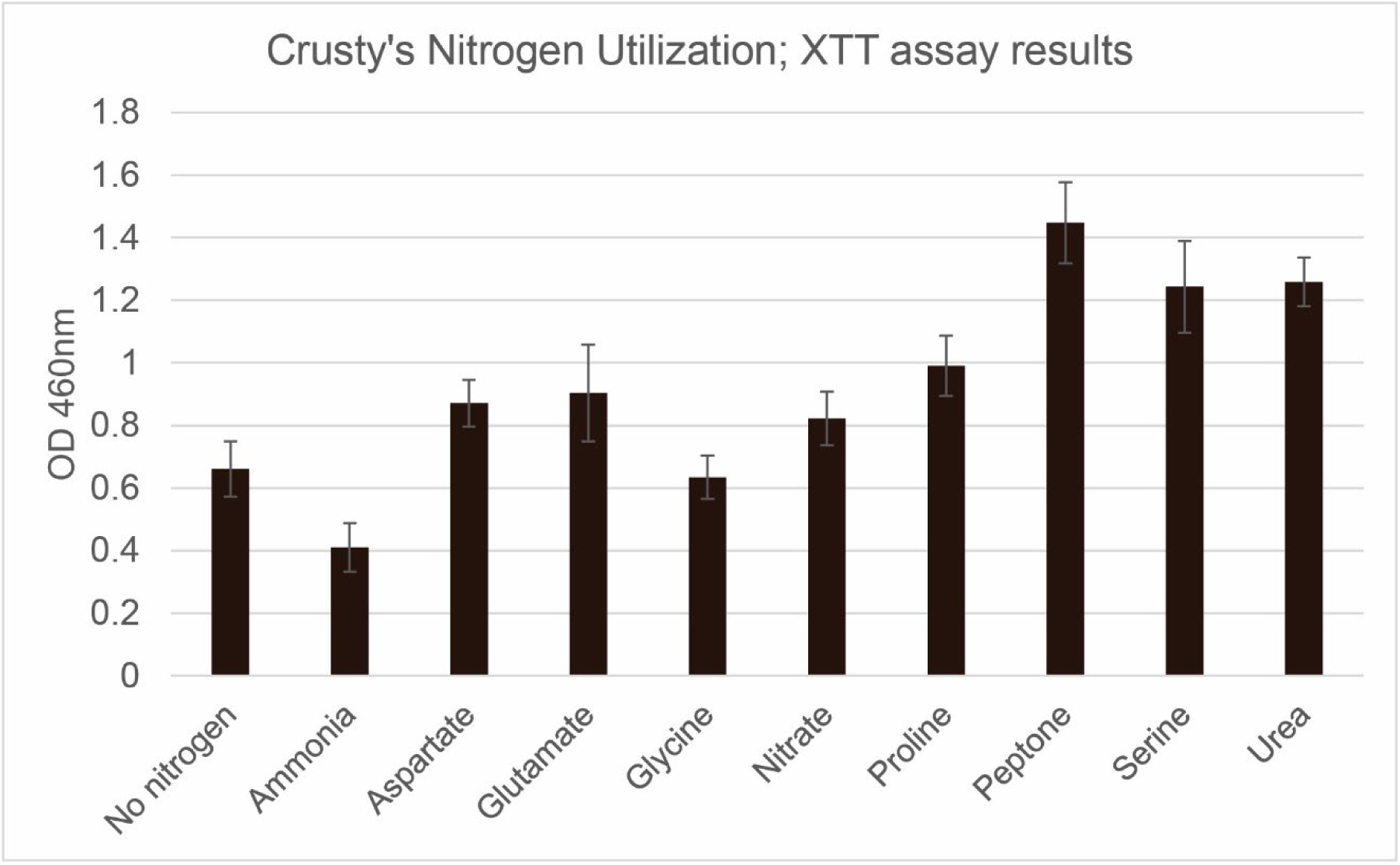
Crusty’s Nitrogen utilization after 10 days of growth, with XTT live growth assay. Peptone showed the highest growth amount, and Ammonia showed the lowest growth amount, oddly lower than no nitrogen source. (n=6, 6 biological replicates)

#### UV-resistance of Crusty

Since Crusty’s colony morphology is drastically dark in color due to melanin deposition in its cell wall, we believed that it would be resistant to high amounts of UV exposure. Performing a serial dilution of Crusty before exposing the cells to the 245nm UV-C light allowed us to ensure that cells were not protected from their 3D clumping nature. In the 0x UV-exposed plate, it seemed that much of the original amount of growth from the control plate was able to grow, accounting for 32% survival by pixel area (**Figure 10**). However, when diluting down to 10x it was obvious that much less of the cells survived with only 8.5% of the control’s area being represented in the UV-exposed plate (**Figure 10**). Finally, on the 100x diluted plate, only 5.4% of the control area was able to grow after UV exposure (**Figure 10**). Considering, *S. cerevisiae* and *E. dermatitidis* are unable to grow at this length of UV-C exposure, Crusty can be considered resistant to UV-C.

**Figure 10:**
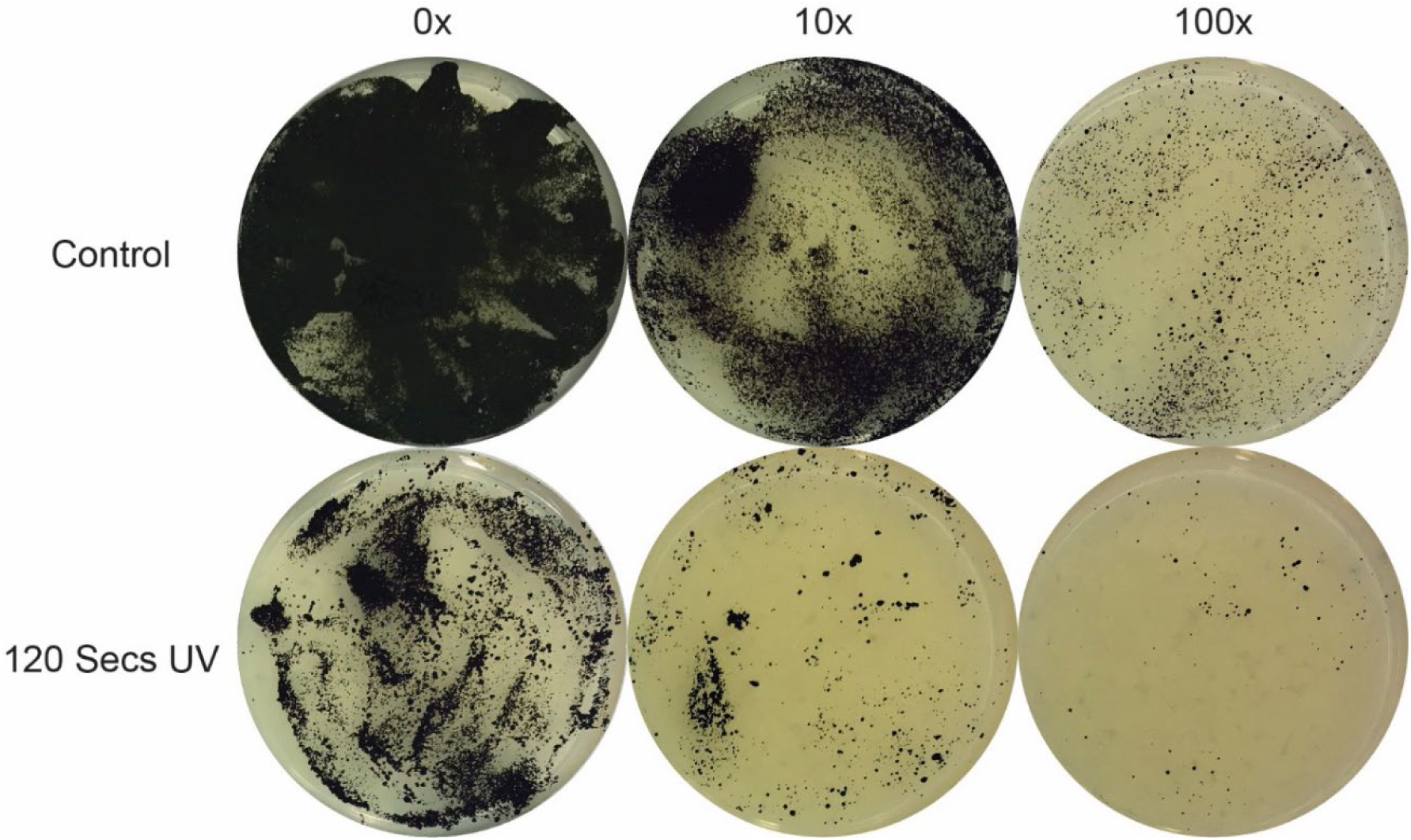
UV resistance of Crusty. Serial dilution of Crusty was performed with 10x intervals, to prevent cells from covering each other and creating an artificially inflated resistance appearance. Control plates had no UV exposure, and UV plates had 120 seconds of 245 nm UV light exposure. As the dilution factor goes up, the amount of cells that survive the UV exposure decrease indicating a coverage protection in the 0x conditions. (n=3, 3 biological replicates)

#### Metal Tolerance of Crusty

Polyextremotolerant fungi are well known for their metal resistance, particularly due to melanin’s metal chelating abilities (Cordero & Casadevall, 2017; Gadd & de Rome, 1988; Hong & Simon, 2007). Therefore, we wanted to determine how resistant Crusty was to various metal species; treatment results were binned based on their inhibition size: none, medium, and high.

Crusty showed no zone of clearing when exposed to 1M FeSO_4_ or 2M LiCl_2_ (**Figure 11** & **Table 3**). Crusty was moderately resistant to 1.5M CuCl_2_, 470 mM AgNO_3_, and CoCl_2_ with a 1.4 cm, 2.8 cm, and 3 cm zones of clearing respectively (**Figure 11** & **Table 3**). Crusty showed the lowest resistance to 10mM CdSO_4_ and 1.5M NiCl_2_ with 4.5 cm and 4.7 zone of clearing respectively (**Figure 11** & **Table 3**).

**Figure 11:**
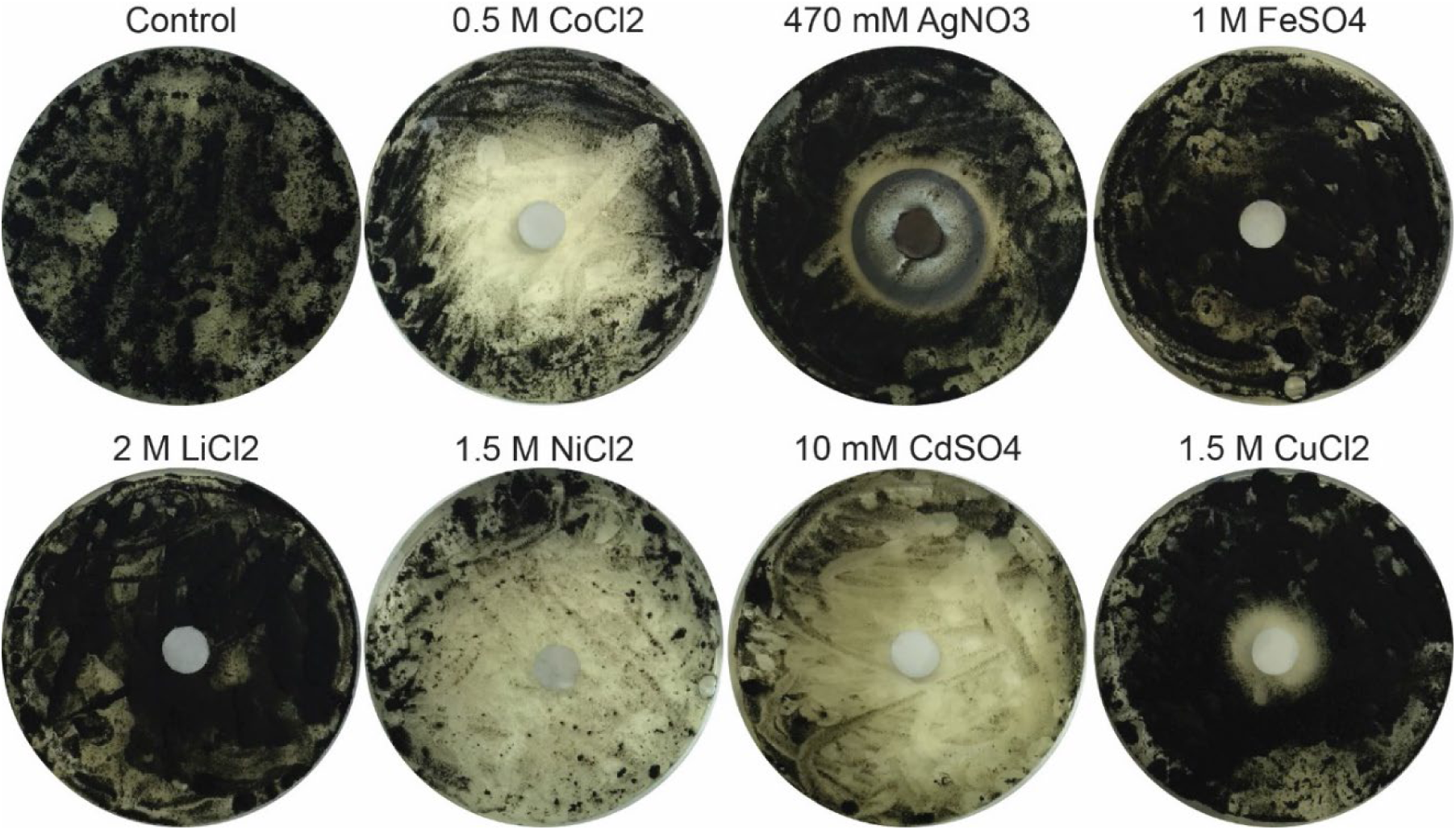
Metal tolerance of Crusty to various metals. In order of most tolerable to least Crusty can completely tolerate FeSO4 and LiCl2 with no zone of clearing, CuCl2 with a 1.4 cm zone of clearing, AgNO3 with a 2.8 cm zone of clearing, CdSO4 with a 4.5 cm zone of clearing, and NiCl2 with a 4.7 cm zone of clearing.

#### Temperature Growth Range of Crusty

To determine the growth range and optimal growth temperature of Crusty we grew Crusty in a 10x serial dilution at five different temperatures from 4 °C to 42 °C. Crusty was capable of slow growth at 4 °C. The optimal growth temperature of Crusty was 23 °C, which showed extensive growth of Crusty at all dilutions (**Figure 12**). Interestingly, at 28 °C Pinky was able to outgrow Crusty making the dilutions above 0x all Pinky colonies with the 0x showing some Crusty colonies with Pinky covering them. Both 37 °C and 42 °C did not promote the growth of Crusty, even though Crusty only remained at those temperatures for 48 hours. Returning Crusty to room temperature after incubation at the higher temperatures did not restore growth, thereby suggesting that exposure to 37 °C and 42 °C might be lethal (**Figure 12**).

**Figure 12:**
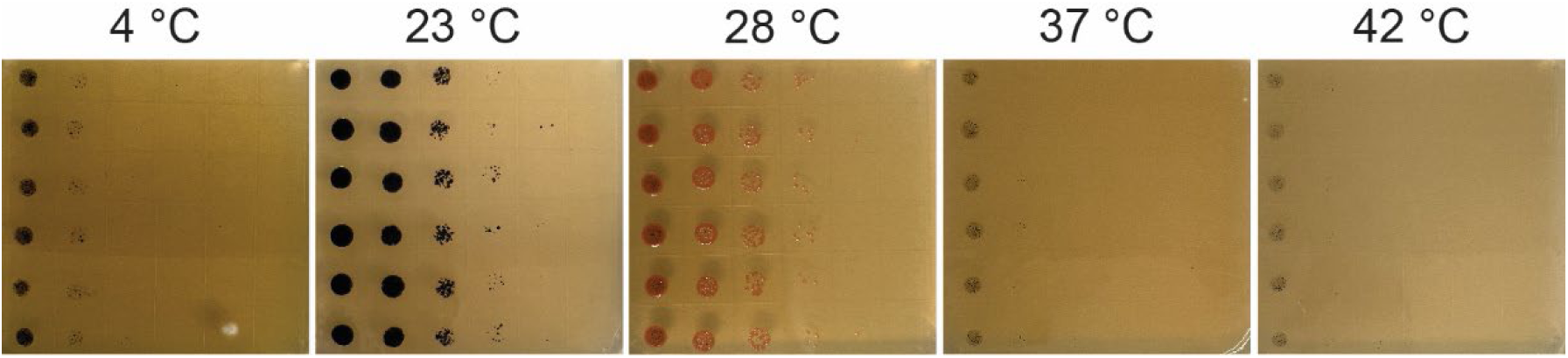
Temperature growth range of Crusty grown on solid media. Serial dilution with a 10x dilution was performed to observe the ability to grow in varying concentrations, and plates were grown for 10 days. Crusty seems to prefer 23 °C for growing but is capable of growing at temperatures as low as 4 °C. Crusty was unable to grow at 37 °C or above, and was killed with 48 hours of exposure at 37 °C or higher.

### Experiments to Decipher the Relationship Between Crusty and its Methylobacterium spp. Symbionts

#### Determination of Endosymbiosis via 16S rDNA Amplification Post-chloramine-T Treatment

Once it was apparent that Crusty had *Methylobacterium* spp. present in its culture, even when they were not forming colonies on a plate, we wanted to determine if the bacteria were inside the fungal cells or not. A method that has been used by previous studies to determine fungal-bacterial endosymbiosis is cell surface sterilization using the Chloramine-T method (Mondo et al., 2012). This is meant to ensure that the only bacteria that DNA is extracted from and therefore 16S rDNA is amplified from, is inside of the fungal cell wall.

We performed this method on Crusty and *S. cerevisiae* as a negative control. When the 16S rDNA region extracted from Crusty prior to Chloramine-T treatment is amplified we see a distinct band, which indicates bacterial presence (**Figure 13**). After Chloramine-T treatment, the 16S rDNA band is still present in Crusty, which indicates the presence of bacteria inside its cell wall. This 16S rDNA band is not present in *S. cerevisiae* which is not known to have a bacterial endosymbiont, and the 16S rDNA band is present for the Light Pinky positive control (**Figure 13**).

**Figure 13:**
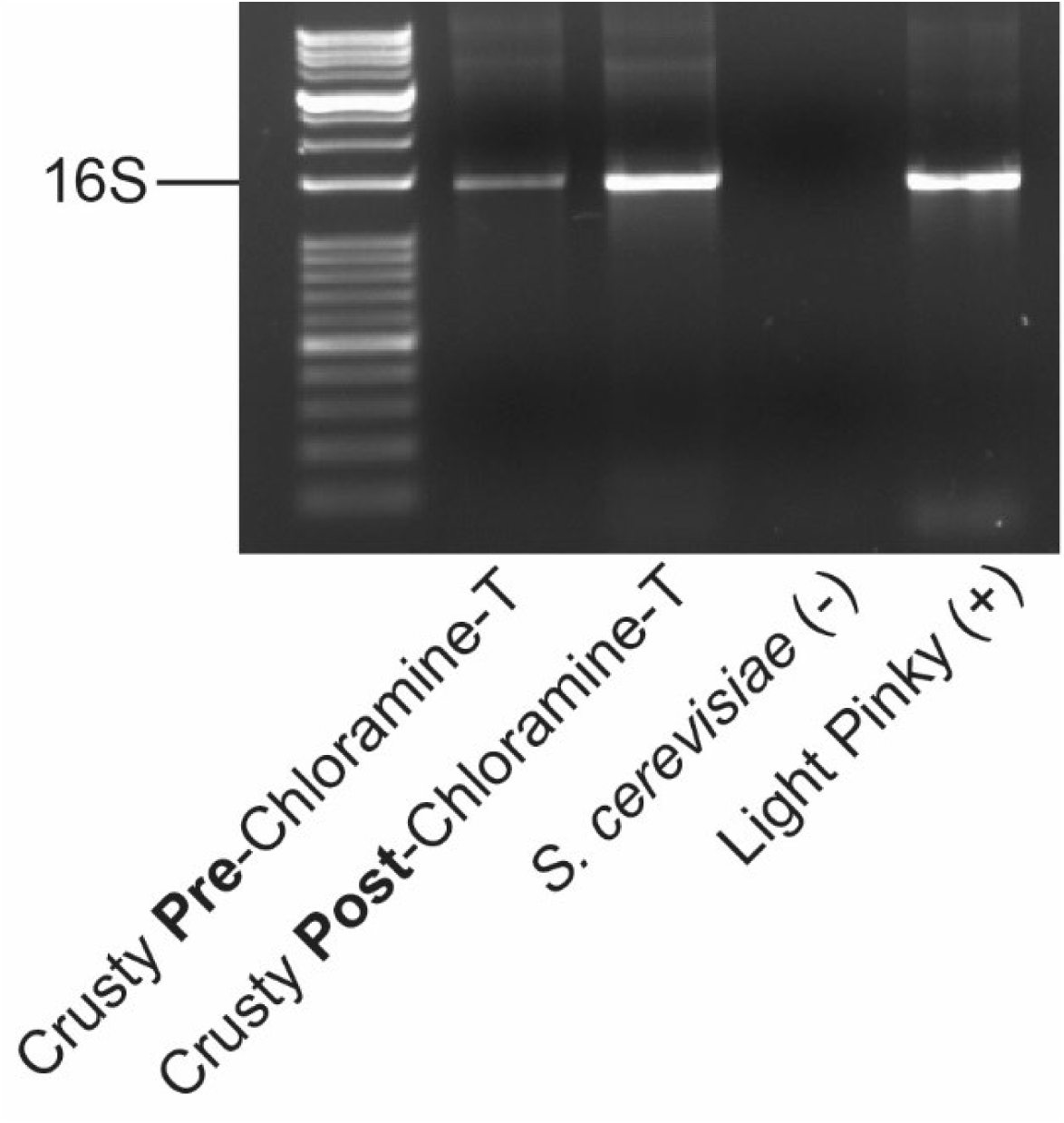
Electrophoresis gel of the PCR amplified 16S region of bacterial DNA. Crusty that was surface sterilized before 16S amplification showed a clear 16S bacterial DNA band, indicating bacterial presence inside of the cells. There was no 16S amplification for the fungus *S. cerevisiae*. There was 16S amplification for the positive control Light Pinky bacteria, and for the non-surface sterilized Crusty.

#### Confocal Microscopy to Observe Bacterial Presence in Fungus

Additional evidence of bacterial endosymbiosis was obtained by performing a bacterial live-dead stain along with a fungal cell wall stain on Crusty. This method was used in another fungal-bacterial endosymbiont study by Partida-Martinez & Hertweck to observe the presence of live bacterial cells within hyphae of Rhizopus (Partida-Martinez & Hertweck, 2005). Replication of their method with the addition of calcofluor white for chitin staining of the fungal cell wall allowed us to observe the relative location of bacteria within Crusty within the same x-axis but this method is unable to resolve a z-axis resolution.

When Crusty cells are stained with PI (red), SYTO9 (green), and calcofluor white (blue) we can see that all three stains will stain parts of the cell (**Figure 14**). The calcofluor white stains only the fungal cell wall, PI stains likely dead bacterial cells, and the SYTO9 stains where presumably live bacterial cells reside. This combination of stains was also performed on the Pinky cells alone, which showed that calcofluor did not stain any cells, and also that not all bacterial cells were capable of being stained with either bacterial stain (**Figure 14**). Additionally, it was obvious that either the bacterial stains were not able to penetrate every fungal cell or that not every fungal cell contained bacteria, because not all fungal cells showed the fluorescence of the bacterial stains (green or red) (**Figure 14**).

**Figure 14:**
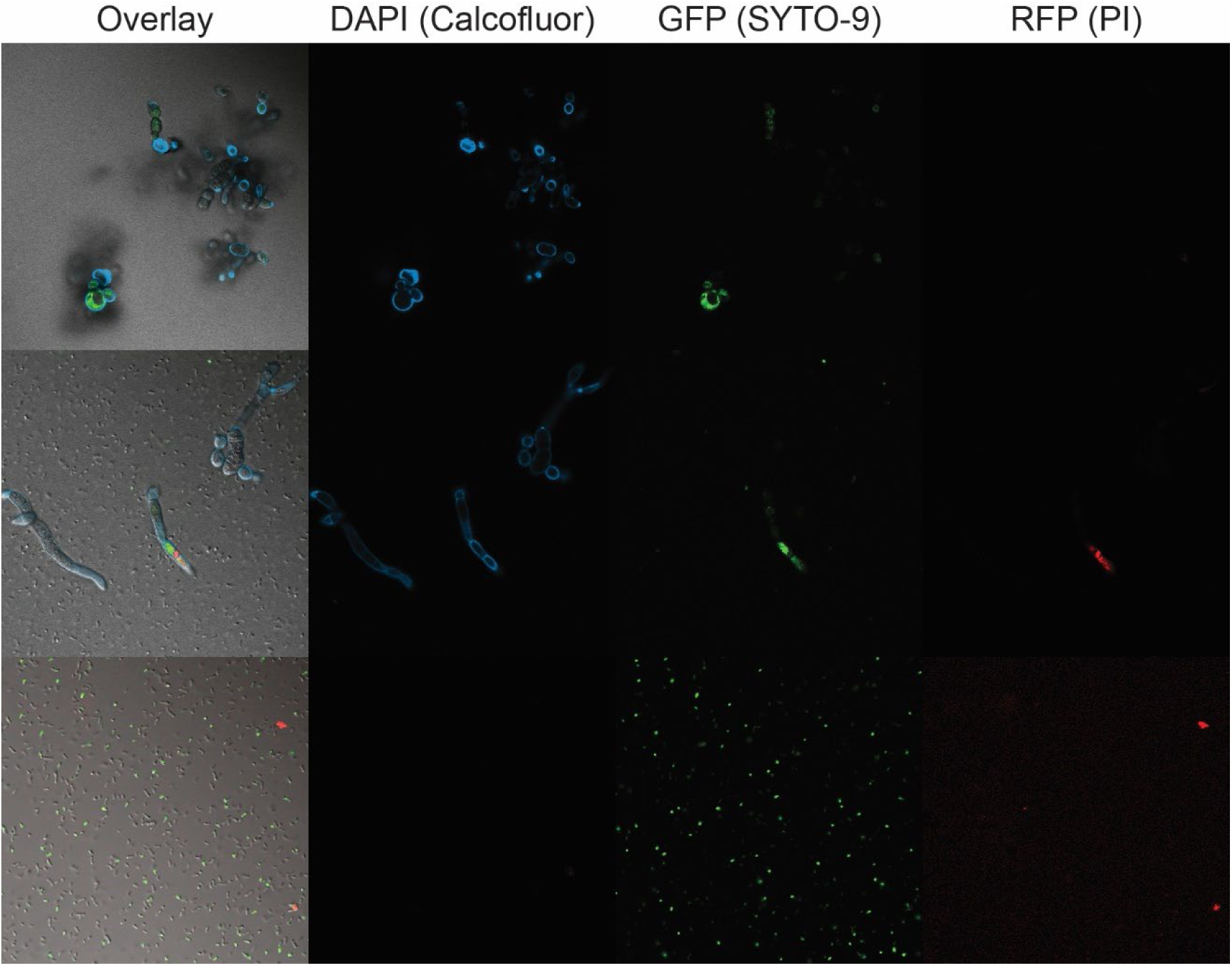
Confocal microscopy of Crusty and Pinky with fluorescent stains. Calcofluor (blue) staining of chitin was done to observe the fungal cell wall which can be seen in the images containing fungi. SYTO9 (green) and PI (red) staining was done to observe the presence of bacteria. SYTO9 was able to stain structures inside of the fungal cells, as was PI. Some bacterial cells were able to be stained with SYTO9 and PI outside of the fungal cells as well.

#### Antibiotics to Treat Crusty of its Methylobacterium Co-occurring Bacteria

In order to create a control strain of Crusty, we attempted to remove its *Methylobacterium* symbionts by using antibiotic exposure. Before we could start with exposing Crusty to antibiotics, we first had to determine which antibiotic would be best to test against Pinky. We performed the Kirby-Bauer antibiotic disk diffusion test using multiple antibiotics individually on a lawn on Pinky (Bauer et al., 1959). The antibiotic that showed the largest zone of clearing was Tetracycline 100 mg/mL (**Figure 15**). However, Tetracycline is only bacteriostatic not bactericidal, meaning it would theoretically not kill and remove Pinky from Crusty but only stop it from growing. The second-best antibiotic that is also bactericidal was Gentamycin 10 mg/mL (**Figure 15**). We decided to use Gentamycin and Tetracycline moving forward in attempting to clear Crusty of its Pinky symbionts.

**Figure 15:**
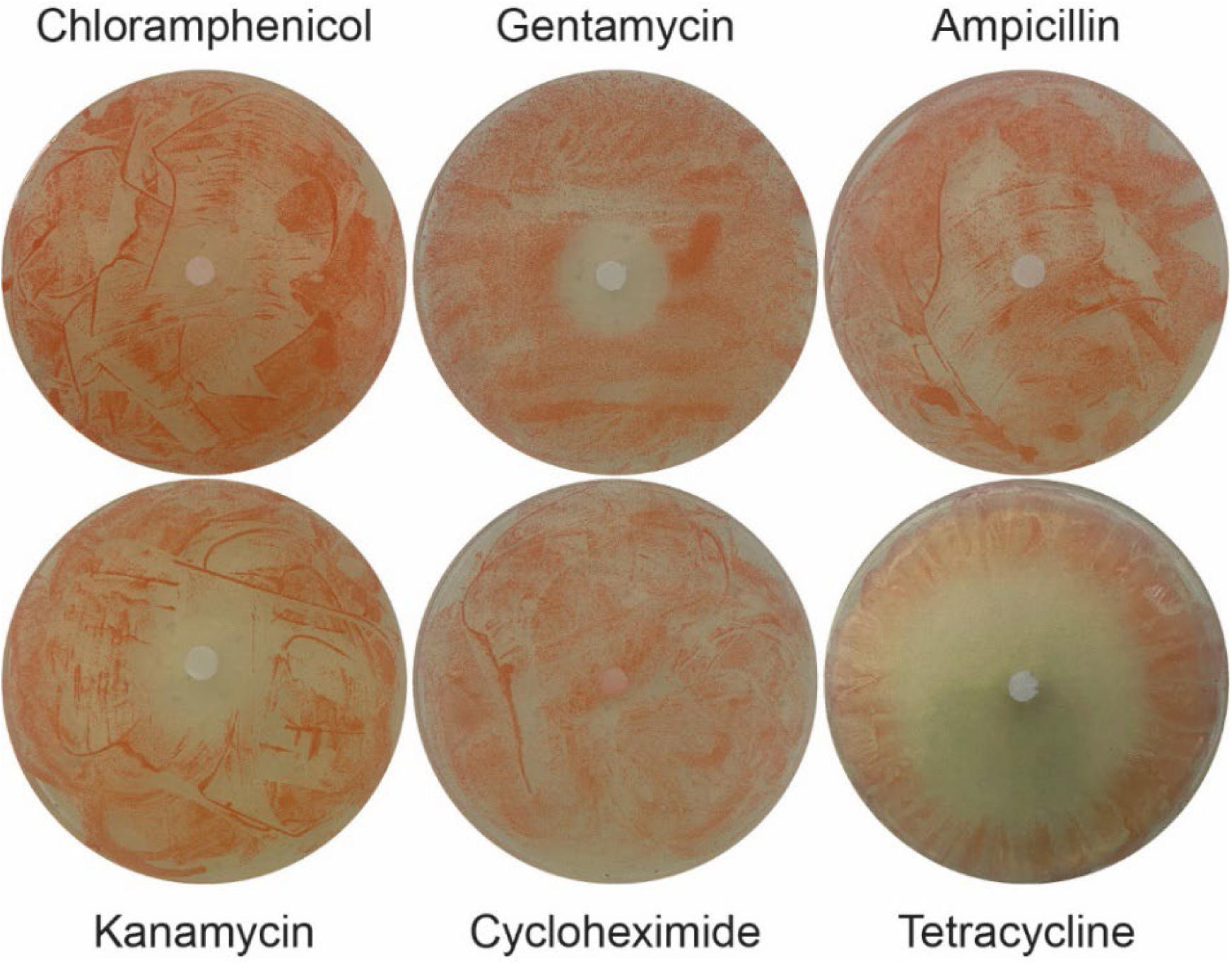
Antibiotic disk test on Pinky to determine best antibiotic to use. Tetracycline showed the largest zone of clearing out of all of the antibiotics, followed by Kanamycin, and Gentamycin. Chloamphenicol, Ampicillin, and Cycloheximide did not create a zone of clearing for Pinky. (n=3, 3 biological replicates)

**Figure 16:**
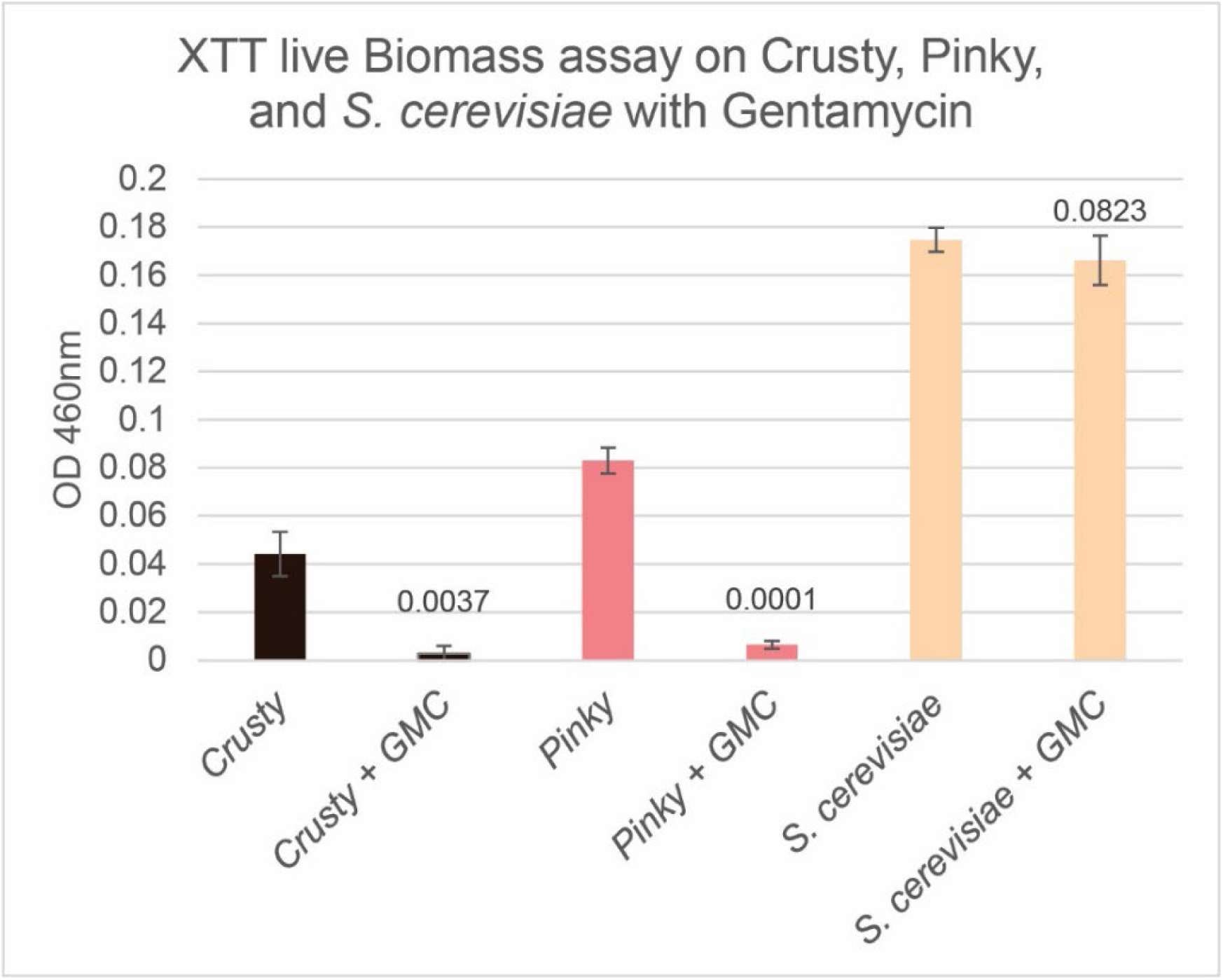
Gentamycin treatment of Crusty, Pinky, and *S. cerevisiae* in liquid culture, with XTT assay live cell confirmation. Crusty and Pinky were both incapable of growth in the presence of Gentamycin, both with p-values less than .01. However, *S. cerevisiae* was capable of growth in the presence of Gentamycin, and the cells exposed to Gentamycin did not have a p-value less than 0.05 compared to the control. (n=3, 3 biological replicates)

#### Gentamycin treatment of Crusty, Pinky, and S. cerevisiae

Using the bactericidal Gentamycin on Crusty as a means of removing its Pinky symbionts did not remove the bacteria. Instead, Crusty was incapable of growth in the presence of this antibiotic. This led us to determine if Gentamycin was capable of killing fungal cells in addition to bacterial cells. Using the XTT assay we tested Gentamycin on Crusty, Pinky, and *S. cerevisiae* to determine if Gentamycin was capable of killing fungal cells and therefore killing Crusty as well, or if it was just killing bacterial cells which, in turn resulted in the inhibition of growth of Crusty. After all three microbial species were grown up in the presence of Gentamycin, only Crusty and Pinky were unable to grow in the presence of Gentamycin. *S. cerevisiae* was capable of growing in Gentamycin’s presence indicating that Gentamycin is not an antifungal (**Figure 15**).

#### Testing Fungal and Bacterial Ability to Grow in the Presence of Tetracycline

Continuing to test antibiotics on Crusty for both the goal of obtaining a bacterial-free culture and now determining how dependent Crusty is on this bacterial presence, we decided to use the more effective bacteriostatic antibiotic Tetracycline. Since it is a bacteriostatic, tetracycline would not be able to entirely kill Pinky but would prevent its growth and therefore stop it from performing metabolic functions that would potentially be essential for its symbiosis with Crusty. We decided to first test Crusty grown in increasing amounts of Tetracycline and then determined the area of growth on plates with and without Tetracycline to observe how Crusty’s growth changes when Pinky’s growth is halted.

When Crusty is grown in increasing amounts of Tetracycline there is a decrease in its ability to grow compared to the carrier 70% ethanol (**Figure 17**). Growing Crusty in 1 μL/mL of Tetracycline all the way up to 4 μL/mL of Tetracycline provides the same amount of growth repression when it is grown on Tetracycline plates, continuing the blocking of Pinky’s growth (**Figure 17**). However, when the same cultures are plated out onto media that does not have Tetracycline added, a dose-dependent response is seen, where the more Tetracycline Crusty was exposed to, the less growth is observed (**Figure 17**). Additionally, no bacteria-free Crusty colonies were able to be recovered from exposure to Tetracycline, even after many attempts.

**Figure 17:**
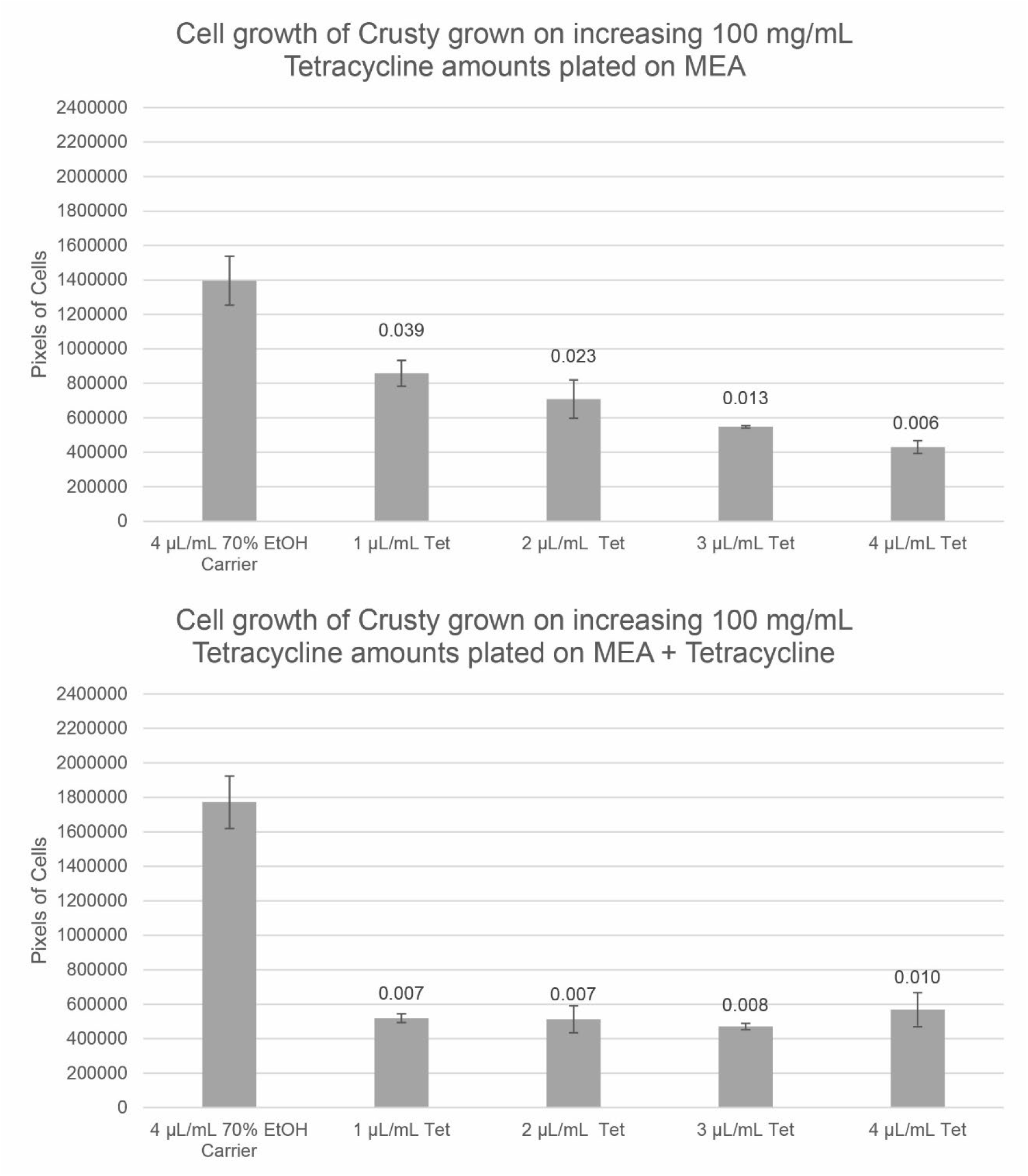
Growth of Crusty on solid media in area of pixels, after various amounts of Tetracycline treatment. After Tetracycline treatment, Crusty that was grown on solid media without Tetracycline added showed an increase in survivability that correlated with a decrease in the Tetracycline treatment, with 4 μL/mL Tetracycline having the least growth p-value 0.006. However, when Tetracycline was added to the solid media that Crusty was grown on after Tetracycline treatment, Crusty had similarly decreased growth at all previous exposure concentrations, and all treatments were significantly different (p < 0.01, n=3, 3 biological replicates).

To ensure that this observation was once again related only to Tetracycline’s effect on the bacteria and not a potential effect on the fungus, we performed the same experiment with *S. cerevisiae* alone and in co-culture with Pinky. This time we chose to use the highest dosage of Tetracycline at 4 μL/mL. When *S. cerevisiae* alone is grown in the presence of Tetracycline its growth actually increases likely due to the presence of the ethanol carrier, indicating no antifungal properties of Tetracycline (**Figure 18**). Additionally, when *S. cerevisiae* is co-cultured with Pinky, there is not a very high decrease in growth (**Figure 18**).

**Figure 18:**
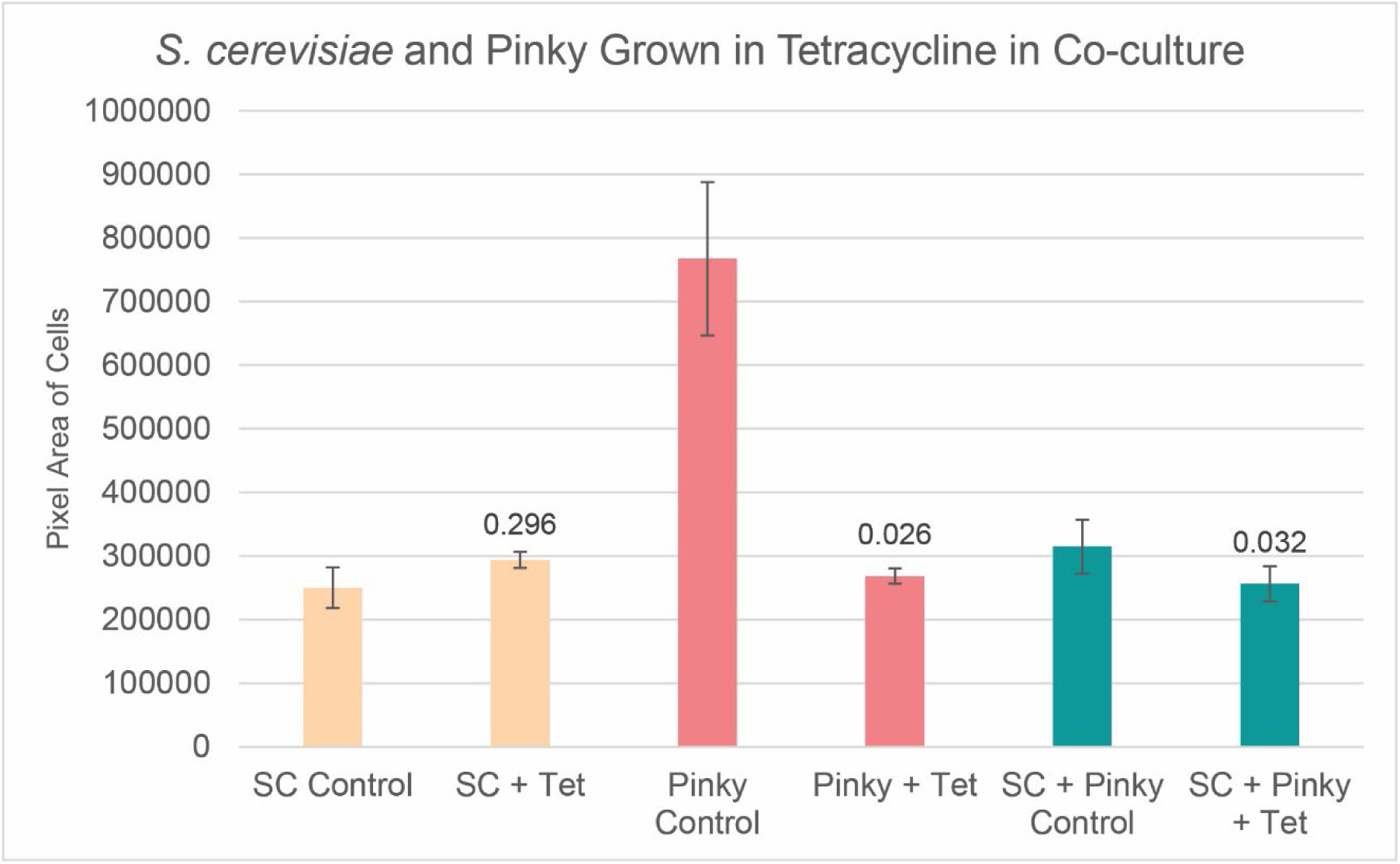
Growth of *S. cerevisiae*, Pinky, and their co-culture after being in the presence of 4 μL/mL of Tetracycline. *S. cerevisiae* had no significant change with the exposure to Tetracycline. Pinky had a significant decrease in growth when exposed to Tetracycline, p < 0.05, n=3, 3 biological replicates. Co-culture of *S. cerevisiae* and Pinky had a significant decrease in growth p < 0.05, n=3, 3 biological replicates.

When comparing the growth inhibition of Crusty, Pinky, and *S. cerevisiae* with and without Tetracycline, we decided to compare the percentage of pixel area between the controls and the 4 μL/mL Tetracycline treatments, due to the differential nature of the colony heights and therefore cell layers unaccounted for by surface area between the three species. When comparing these microbes’ percentage of growth when grown in 4 μL/mL of Tetracycline compared with the ethanol carrier control, Crusty experienced the most drastic decrease in growth, where only 31% of cells were able to grow in the presence of Tetracycline compared to the control growth (**Figure 19**). Pinky experienced the second least growth at 35% compared to its control, and *S. cerevisiae* in co-culture with Pinky only had 81% growth compared to its control (**Figure 19**). This difference in cell area is very apparent when observing the plates themselves, where the *S. cerevisiae* control plates looks almost identical to the 4 μL/mL of Tetracycline plate, whereas both the Crusty and Pinky plates show a large decrease in colonies on the Tetracycline plates compared to their controls (**Figure 20**).

**Figure 19:**
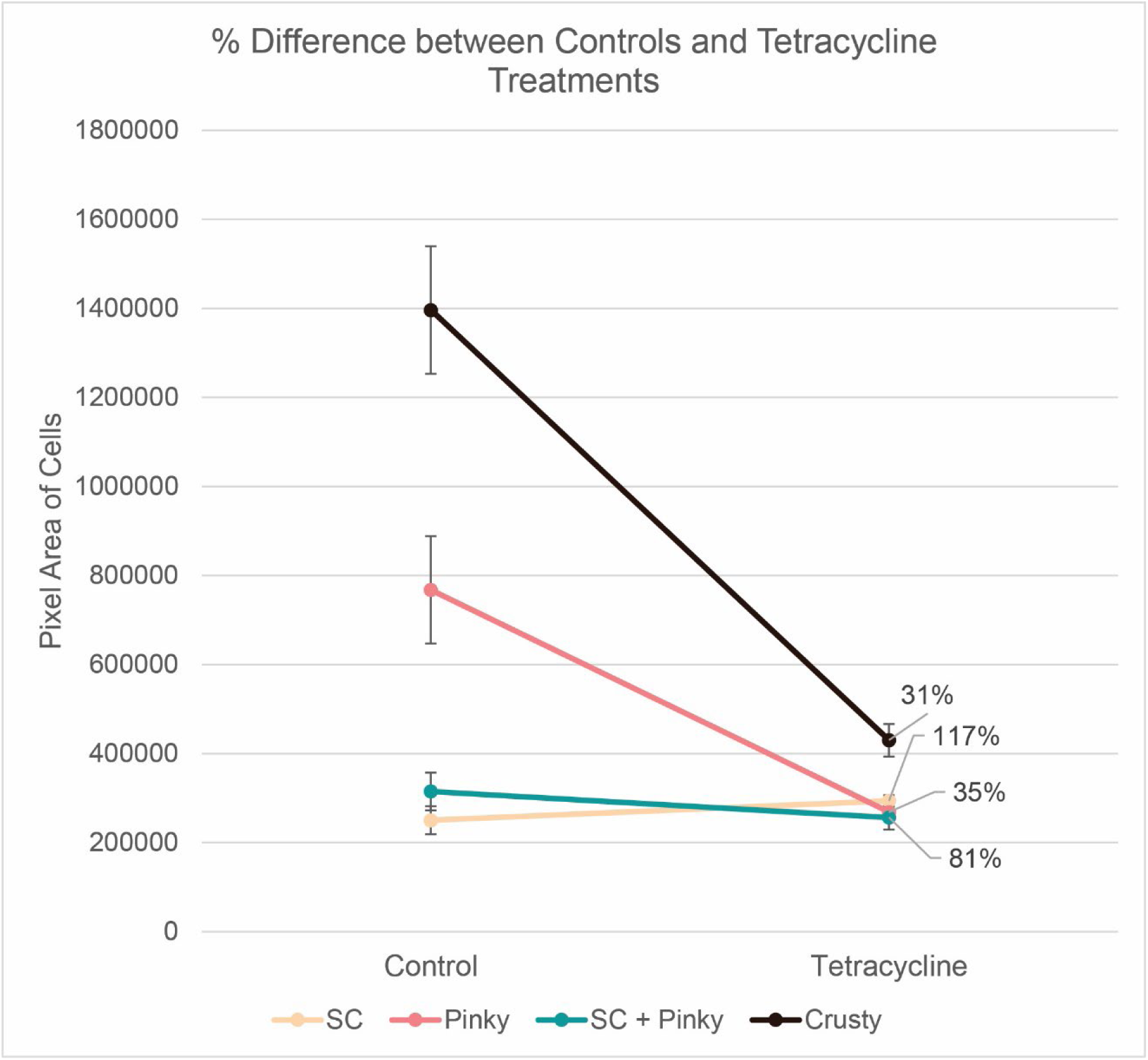
Percent difference between the Pixel area of *S. cerevisiae*, Pinky, Crusty, and the co-culture of Pinky and *S. cerevisiae* grown with and without 4 μL/mL of Tetracycline. Only 31% of control Crusty was able to grow compared to growth in the Presence of Tetracycline. 117% of S. cerevisiae was able to growth in the presence of Tetracycline compared to its control growth. Pinky had a similar growth decrease as Crusty, in that only 35% of the original growth amount was seen in the presence of Tetracycline. Finally, 81% of cellular growth was seen in the S. cerevisiae-Pinky co-culture in the presence of Tetracycline compared to the control.

**Figure 20:**
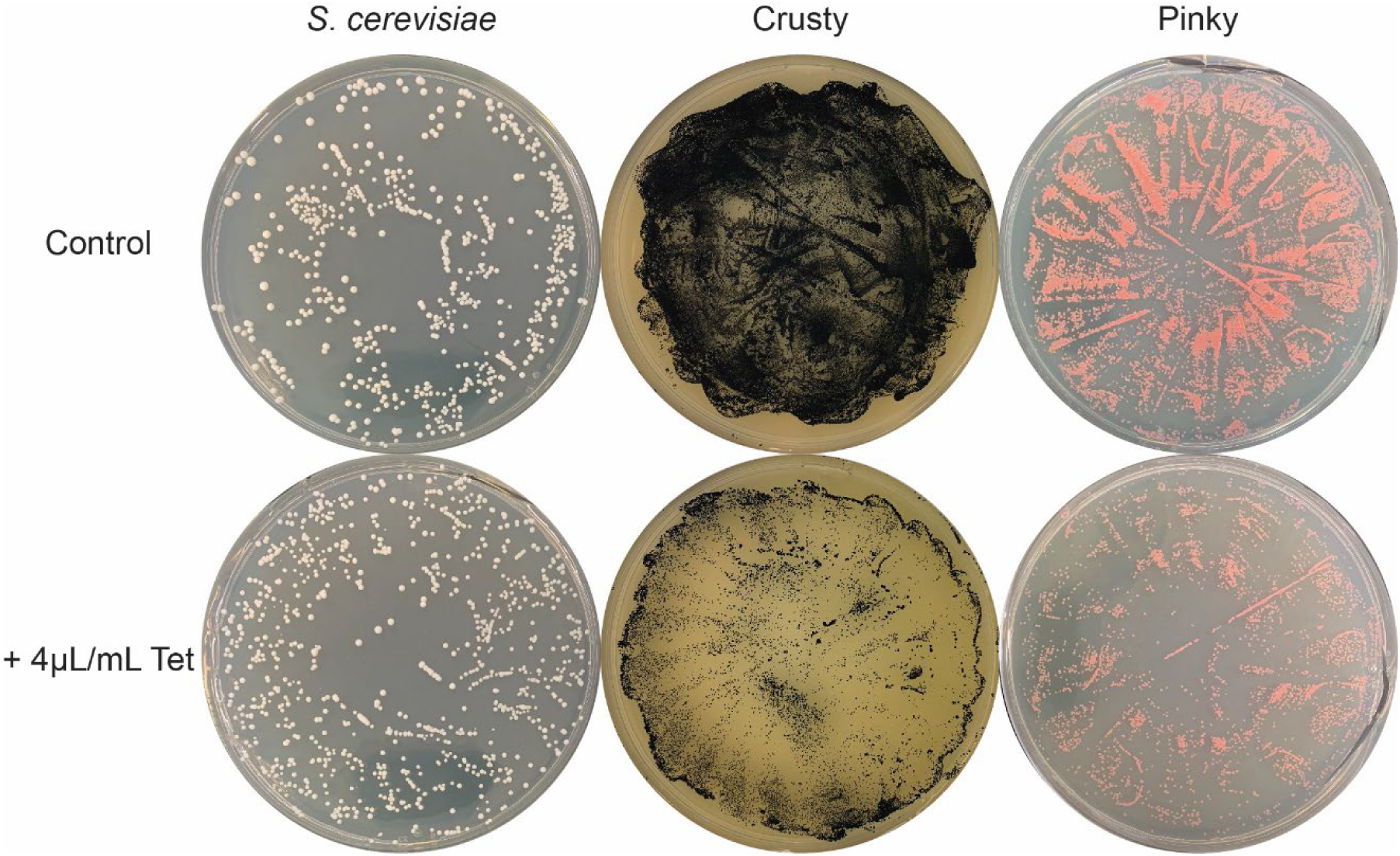
Representatives of the CFU plates used to determine pixel area of *S. cerevisiae*, Crusty, and Pinky control and after 4 μL/mL Tetracycline exposure. The difference in area covered in cells on the control plates for Crusty and Pinky compared to the 4 μL/mL of Tetracycline plates shows a great decrease in survivability or growth.

### Microbial Interaction Experiments

#### Tri-culture Experiments in Carbonless BBM to Observe Nutrient Acquisition from C. sorokiniana to Crusty

Our larger question in the isolation and identification of multiple polyextremotolerant fungi has been to determine if they can undergo lichen-like interactions with photosynthetic algae and nitrogen-fixing bacteria. When this experiment began, we did not know if the *Methylobacterium* spp. harbored by Crusty contained nitrogen-fixing metabolism or not. However, we assumed that they would, based on their phylogenetic relationship with Rhizobiales and other nitrogen fixing *Methylobacterium* spp. Therefore, we thought that the Crusty-Pinky symbiosis would be the more successful fungus to try in attempting a lichen-like co-culture (or tri-culture) with the algae *C. sorokiniana*, in addition to the well-known production of growth hormones by *Methylobacterium* spp. which would likely assist in this tri-culture. The algae *C. sorokiniana* was also chosen due to its phylogenetic relationship to lichen forming Trebouxioid algae, and convenience due to its availability in our stock collection. We performed a tri-culture, and respective mono- and co-cultures for controls, of Crusty, Pinky, and *C. sorokiniana* UTEX 1230 in BBM with no carbon source and performed the XTT assay to observe if Crusty had increased active metabolism (the direct measurement of the XTT assay) with only an algae’s secretion of sugar as a potential source of carbon after 27 days.

Tri-culture of Crusty with Pinky and *C. sorokiniana* did not result in increased metabolism, compared to Crusty without any added microbes (**Figure 21**). Additionally, Crusty grown in the presence of *C. sorokiniana* only (with its presumed Pinky endosymbiont) also did not have increased metabolism compared to Crusty without any added microbes (**Figure 21**). The only conditions that did increase the active metabolism of Crusty was when 10 μL instead of 100 μL of Pinky was externally added to the tri-culture (**Figure 21**). Most notably, Crusty was capable of active metabolism in a carbonless medium without a photosynthetic microbe present, only relying on itself and Pinky, which was not initially thought to provide a carbon source to Crusty (**Figure 21**).

**Figure 21:**
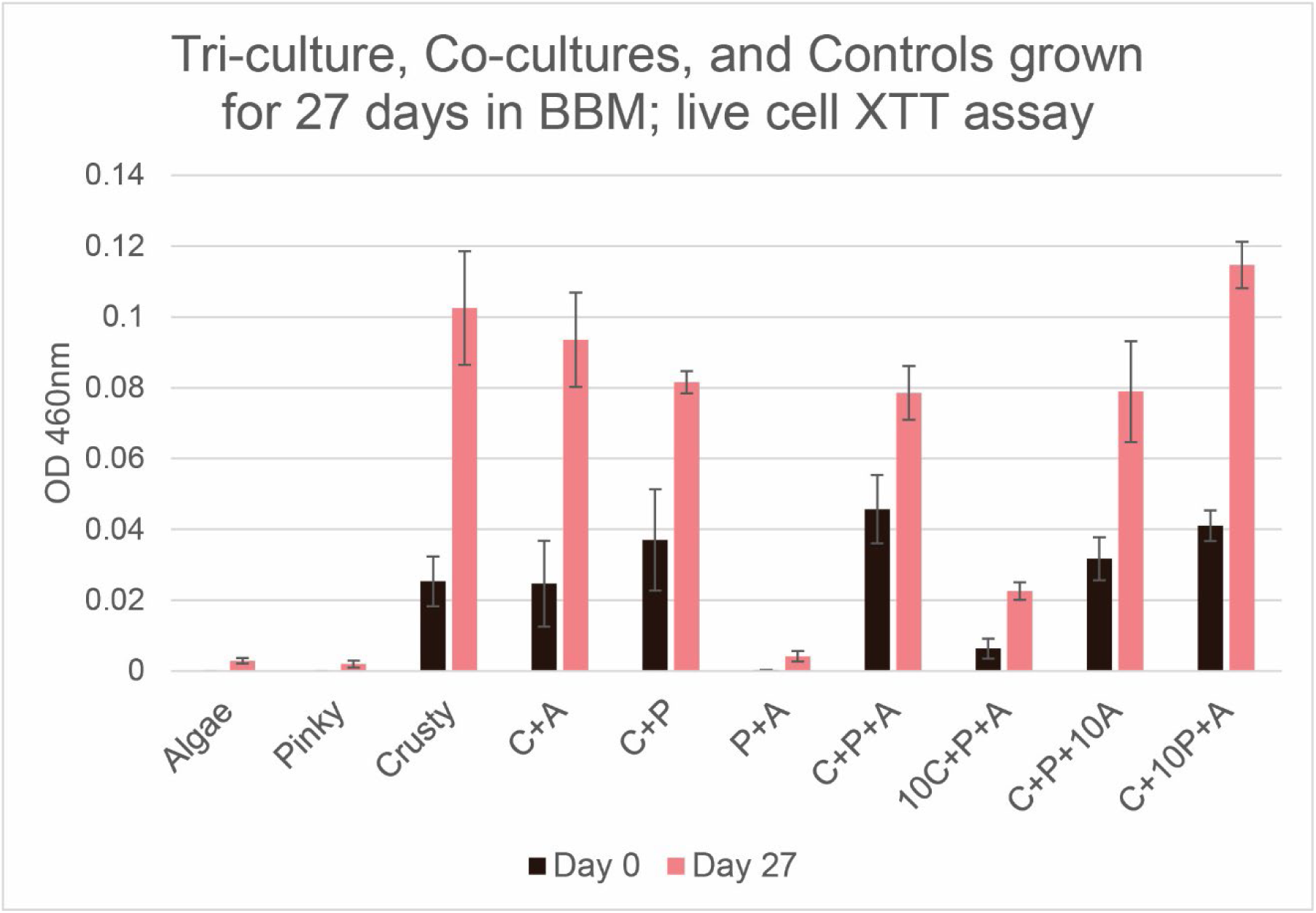
XTT live cell assay OD showing fungal cell growth. All microbes were added to the flask in 100 μL amounts, except the conditions that have 10 before their letter which indicates only 10 μL of that microbe was added. The Algae (*C. sorokiniana*) and Pinky were unable to be properly detected via the XTT assay, which was optimized for Crusty. Crusty growth in BBM without added external microbes had the second highest growth amount after Crusty + 10 μL of Pinky + Algae, but the difference between these is not significant. None of the other conditions showed a significant increase or decrease in growth of Crusty, except when 1/10^th^ the volume of Crusty was added initially (10C+P+A). (n=3, 3 biological replicates)

#### Crusty-Pinky +/- Carbon, Nitrogen, Light

Due to the interesting observations seen in the tri-culture experiment, where Crusty was capable of active metabolism in a carbonless environment, we decided to re-test the Crusty-Pinky symbiosis against multiple media drop-out conditions. We grew Crusty-Pinky in BBM media that either did or did not contain mannitol as a carbon source or nitrate as a nitrogen source. All conditions were either exposed to a 12-hour light/dark cycle or kept permanently in the dark (n=3, 3 biological replicates). This experiment was done to determine whether the Crusty-Pinky symbiosis was capable of active metabolism in the absence of essential carbon and nitrogen sources, and whether light induced active metabolism in these conditions. If either or both organisms were capable of active metabolism without any external carbon or nitrogen source, then it is likely that Pinky could be producing a carbon or nitrogen source for Crusty, or that Pinky is engaging in anoxygenic photosynthesis.

When Crusty-Pinky was grown up in BBM with no carbon, no nitrogen, and no light, the XTT assay showed that active metabolism decreased from day 7 to day 14 (**Figure 22**). However, the CFU assay for these same conditions showed on day 7 a pure culture of Crusty with no Pinky escaping the Crusty cells, but on day 14 there was a mixed culture of Crusty and Pinky colonies (**Figure 23**). Interestingly, when light is introduced to the no nitrogen and no carbon conditions (-N -C), there is a significant increase in active metabolism from day 7 to day 14 (**Figure 22**). Although, this does not reflect an increase in cellular growth as seen in the CFU assay. The CFU plates for Day 7 and 14 of the no nitrogen, no carbon, plus light contained the same colony number as the no light plates (**Figure 23**). This trend of an increase in active metabolism with the addition of light to the various conditions tested, is seen in every other condition tested (+N, +C, +N +C), and is statistically significant with p < 0.05 (n=3) in all except the +N +C conditions (**Figure 22**).

**Figure 22:**
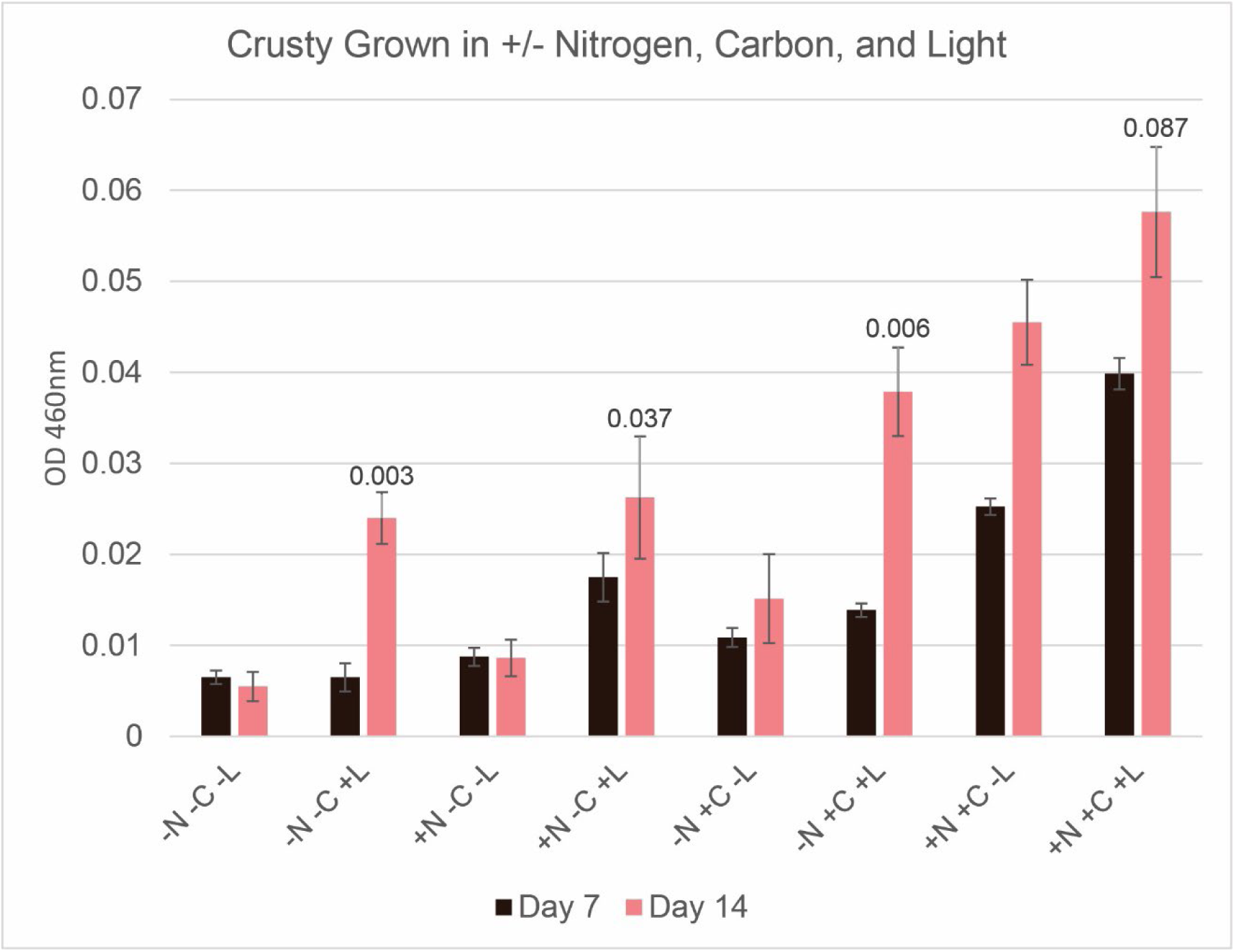
XTT assay results of Crusty growth in BBM with and without the addition of a carbon source, and nitrogen source, and 12 hours of light per day. Crusty was capable of growth without a carbon source or a nitrogen source, as long as it had 12 hours of light per day, p = 0.003. When an external nitrogen source was added growth did not increase if light was not available, but growth significantly increased when light was available compared to no light, p = 0.037. When a carbon source was added growth was capable even without light, but was much higher with light present, p = 0.006. When all nutrients were provided, growth was capable even without light, but with added light growth was not significantly higher than without light, p = 0.087. (n=3. 3 biological replicates)

**Figure 23:**
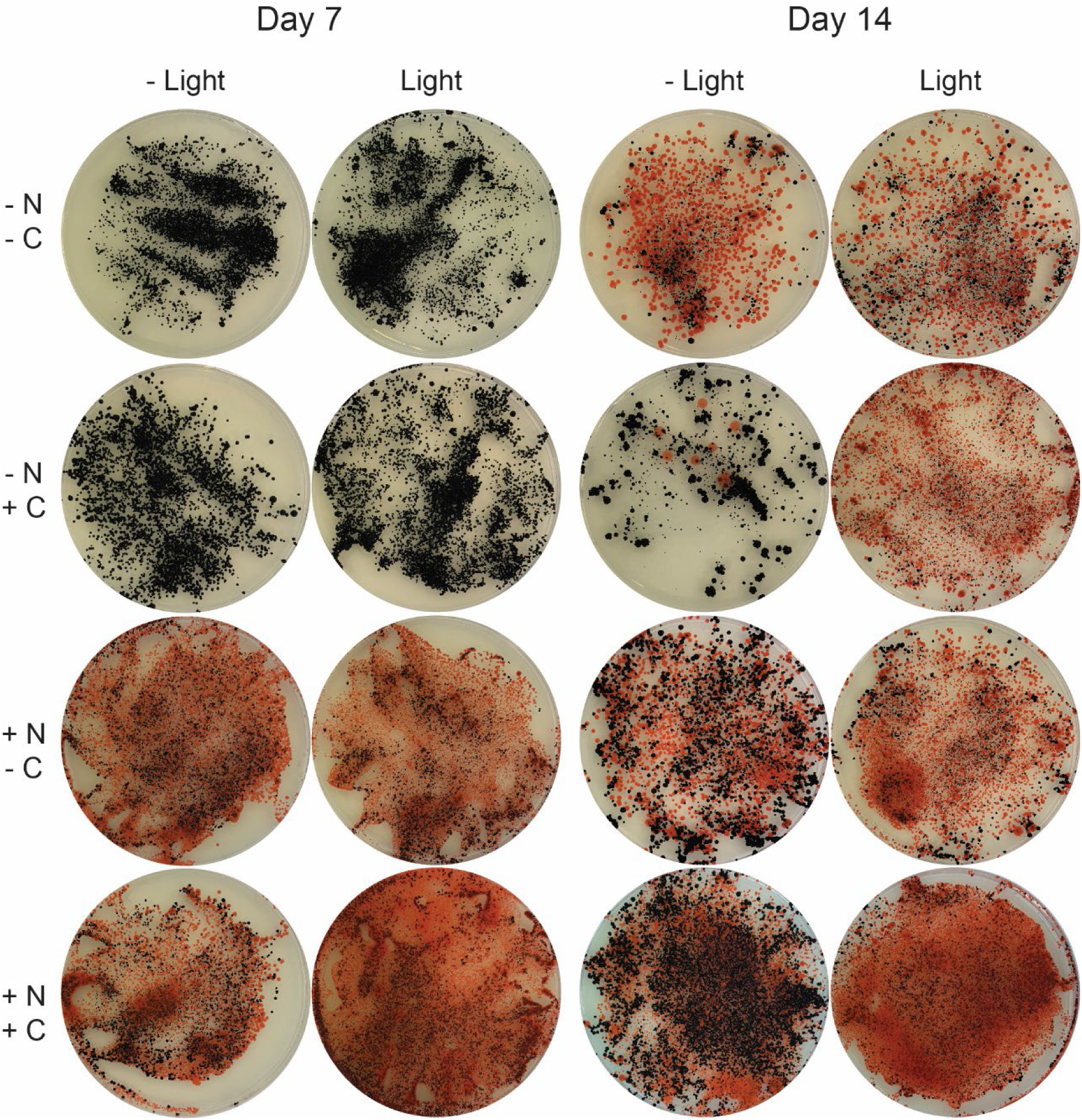
CFU plates of Crusty growth with and without carbon, nitrogen, and light at day 7 and day 14. On day 7 Crusty grown without a nitrogen source prevented Pinky from coming out of its cells, with or without light. However, when a nitrogen source was present, Pinky was able to escape Crusty by day 7. On day 14 only the no nitrogen, added carbon, no light had limited Pinky release from Crusty cells. All other conditions allowed Pinky to leave the cells. Although, if light was not present the amount of Pinky colonies seen outside of Crusty were much less than if light was present.

CFU plates of each condition show a rough ratio of Crusty to “escaped” Pinky that would be reflective in the XTT assay performed. Interestingly, addition of carbon leads to the retention of Pinky on day 7 whether there is light exposure or not (**Figure 23**), and on day 14 only very few Pinky colonies can be seen in the no light but added carbon condition (**Figure 23**). However, when light exposure is present, Pinky is capable of being released in the carbon conditions on day 14 (**Figure 23**). The rest of the CFU plates whether light, carbon, and/or nitrogen are present show a mixture of Crusty and Pinky colonies on all plates (**Figure 23**).

#### Overlay of Algae over Methylobacterium spp. Spots

The reason for why Pinky is associated with Crusty is still largely unknown. However, *Methylobacterium* spp. are known to produce plant phytohormones (Koenig et al., 2002; Tani et al., 2012). Therefore, to determine if Light Pinky and Dark Pinky were capable of growth stimulation of algae, we decided to observe how the individual Pinky’s would interact with the algae *C. sorokiniana*. All *Methylobacterium* isolates and the combination of Light and Dark Pinky were spotted onto MEA, which is a medium that they all readily grow on. Then, we overlayed either 0x, 10x, 100x, or 1000x diluted *C. sorokiniana* in 0.5% agar BBM. This allowed us to observe how the bacteria on the spots would affect the algae directly above them.

Growth stimulation of *C. sorokiniana* when overlayed on top of Pinky (both Light Pinky and Dark Pinky) at every dilution seemed minor (**Figure 24**). There was only a slight growth increase directly over the bacterial spots, but the algae grew in an even coating across the plates. However, when Light Pinky and Dark Pinky were spotted individually, both were capable of stimulating algae growth directly above their spots (**Figure 24**). Light Pinky was more capable of enhancing the growth of the algae than Dark Pinky because there was more algal growth on the algae that was directly touching the spots over Light Pinky than over Dark Pinky. Interestingly, both Dark Pinky and Light Pinky also show a slight zone of inhibition at the highest concentration of *C. sorokiniana* (0x). The area directly around the middle dot surrounded by the other eight dots is paler for both plates (**Figure 24**). Directly over the dots still shows growth enhancement, as there is also growth enhancement over each of the dilutions for Light and Dark Pinky. Additionally, both Light Pinky and Dark Pinky seem to enhance colony formation of *C. sorokiniana* versus planktonic growth when compared to their combination (Pinky) or the other *Methylobacterium* spp. P3, on the other hand, had a complete zone of inhibition, clearing out any algal growth especially on the 10x diluted sample (**Figure 24**). In the 0x sample, it appears that there is some growth stimulation on the corner bacterial dots though. However, the zone of inhibition becomes larger as the algal concentration decreases to where there is almost no algal growth in the 1000x sample (**Figure 24**).

**Figure 24:**
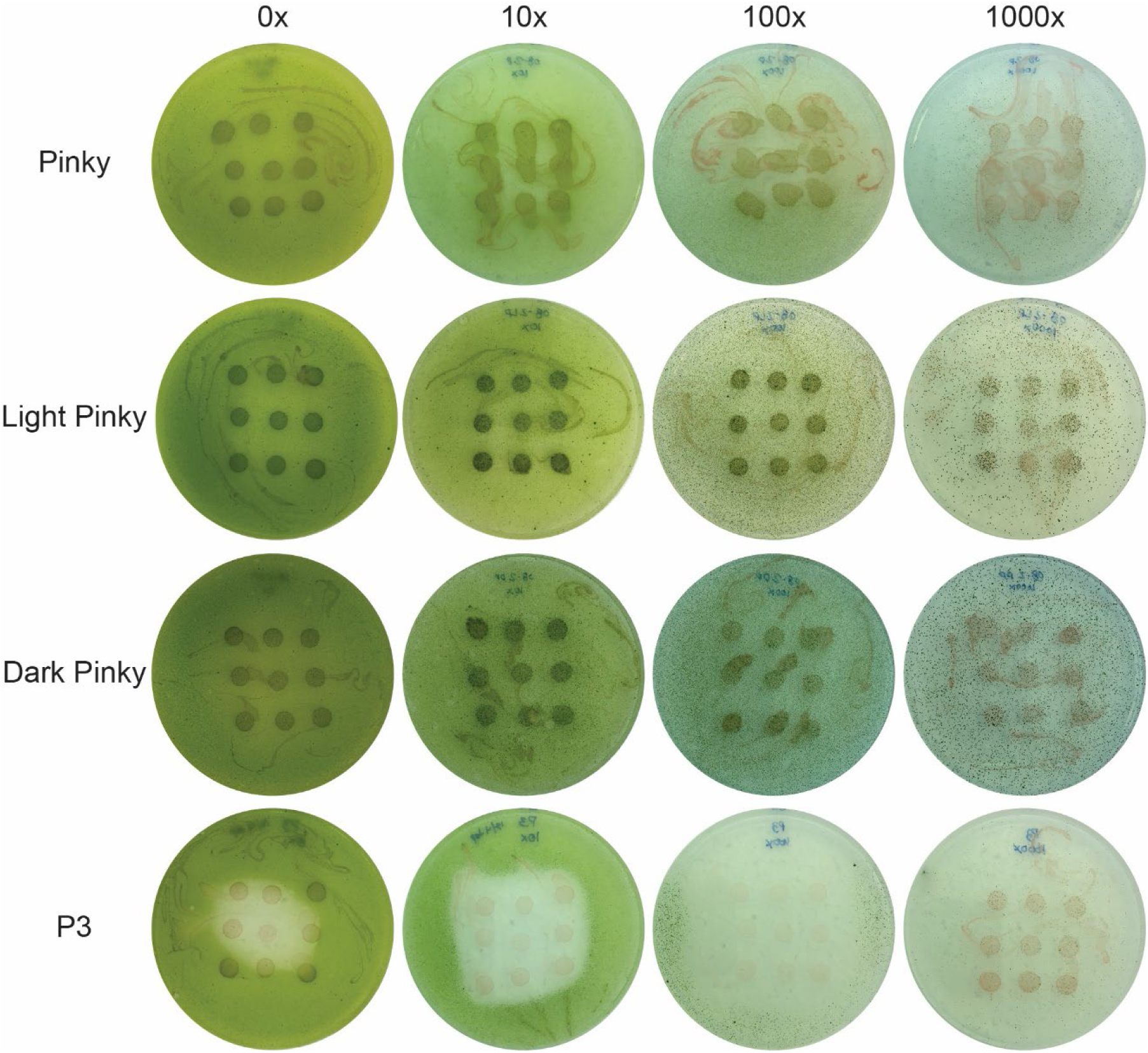
*C. sorokiniana* overlay on top of bacterial spots to determine in-direct growth promotion of algae from bacteria. Pinky, the combination of both Light Pinky and Dark Pinky, showed a slight growth increase of the algae that were directly on top of the bacterial spots. Light Pinky and Dark Pinky both showed very robust increase in algal growth at all dilutions of the algae. Additionally, they seemed to enhance the colony-forming nature of the algae, as it formed spots on the agar instead of a lawn. They also showed some slight growth decreasing ability in the 0x algae growth in the area directly around where the bacterial spots were placed. P3 on the other hand showed complete zones of inhibition, directly killing or repressing the growth of the algae that were in the vicinity of its colonies. (n=3, 3 biological replicates)

## Discussion

### Crusty is a Novel Polyextremotolerant Fungi

In this study, we have described a novel polyextremotolerant fungal species that appears to harbor endosymbiotic bacteria. The morphological group of polyextremotolerant fungi is not historically a widely studied group of fungi, yet active research into this group has recently increased which will likely lead to the discovery and characterization of new polyextremotolerant fungi in the near future. Crusty is likely a member of the genus *Neophaeococcomyces* based on the current ITS sequences and phylogenetic analyses performed. However, documentation of this genus is limited to sequences in the NCBI database, with no full publications of experiments on members of the genus itself, and only the initial descriptions required for novel species identification (Crous et al., 2015; Moreno-Rico et al., 2014; Réblová et al., 2016). With this detailed description and experimentation of the novel species Crusty we expect to gain additional knowledge into this cryptic genus.

### Mimicking the Biological Soil Crust Symbiosis with Crusty, Pinky, and C. sorokiniana

Biological soil crusts are vastly complex biofilms, containing millions of different taxa with a range of ecological relevance (Belnap et al., 2001; Belnap & Lange, 2003). Therefore, studying them for their microbial interactions can be quite difficult, as microbial interactions in any microbiome are very complex. In an attempt to simplify this complex consortium, we whittled this down to what we believe to be some of the most important niches within a biological soil crust: a nitrogen source provider, a carbon source provider, and a structural biomass adherence and/or provision of abiotic stress tolerance (melanin). The first two niches are metabolically essential, in that there is no external source of nitrogen or carbon for the biological soil crust biofilm (Belnap, 2002; Belnap & Büdel, 2016; Li et al., 2012). Both niches are largely held in the beginning by cyanobacteria, particularly *Microcoleus* spp. (Belnap, 2003), then algal and nitrogen-fixing bacteria participate in those roles during the later stages of biological soil crust development (Bowker et al., 2002; Lan et al., 2012). Since we had a fungus with symbiotic bacteria that we believed at the time to have nitrogen assimilating capabilities and an alga of the family Trebouxiophyceae, a common lichen photobiont (Ahmadjian, 1982; Kroken & Taylor, 2000; Triebel & Dagmar, 1992), we decided that these would be our best candidates for attempting a simulated biological soil crust symbiosis experiment, the tri-culture experiment.

The algae *C. sorokiniana* was chosen initially because it belongs to one of the most extensive lichen photobiont groups Trebouxiophyceae. However, we did not know at the time that *C. sorokiniana* does not perform the same metabolic functions that its lichenized Trebouxioid relatives have, mainly the production and secretion of polyols, particularly ribitol, as a carbon source (Richardson & Smith, 1968; Richardson et al., 1967). Instead, *C. sorokiniana* and other *Chlorella* species do not secrete any polyols (Gustavs et al., 2011), they only produce and excrete glucose and maltose (Fischer et al., 1989; Kessler et al., 1991). While these are still carbon sources that can be used by any microbes in its vicinity, they may not be a good analogue for the algal portion of the biological soil crust symbiosis that we had hoped to mimic. Mainly, glucose as a carbon source induces carbon catabolite repression, which is a major metabolic and transcriptional regulator for many other “non-ideal” nutrient metabolite processes such as nitrogen metabolism (Dowzer & Kelly, 1991; Nair & Sarma, 2021; Portnoy et al., 2011; Ries et al., 2016). The reason we did not see a distinct increase in the growth of Crusty when co-cultured or tri-cultured with *C. sorokiniana*, may have been because of this distinct metabolic and therefore interactive difference.

We observed an increase in growth in one of the tri-culture flasks containing 10 μL of added external Pinky with 100 μL of *C. sorokiniana* and Crusty individually. These conditions had the highest increase in active metabolism across the tri-cultures, including higher metabolism than cultures inoculated with all three microbes in equal amounts. An explanation for this, could lie in the bacterial component. *Methylobacterium* spp. are known to produce plant phytohormones, which increase the growth of the plants in exchange for methanol which is released from plant stomata (Kutschera, 2007; Trotsenko et al., 2001). The *Methylobacterium* produce multiple types of these compounds, mainly cytokinins and indole acetic acid (Ivanova et al., 2001; Koenig et al., 2002; Meena et al., 2012), which have been shown to increase the growth of algae (Khasin et al., 2018) and increase fungal growth (Fu et al., 2015; Kulkarni et al., 2013; Sun et al., 2014). These phytohormones are shown to have dose-dependent response in these systems, only stimulating growth at low levels, with higher levels causing growth repression (Kulkarni et al., 2013; Sun et al., 2014). It could be that when Pinky was added at 1/10^th^ of the ratio of the algal and fungal components to the tri-culture system, it did not exceed the amount of hormone production for growth promotion. Additionally, less bacterial initial inoculum possibly prevented the bacteria from scavenging more of the glucose carbon source from Crusty.

We were able to confirm that both Light Pinky and Dark Pinky have growth promoting capabilities with the *C. sorokiniana* plate overlay experiments, the growth of *C. sorokiniana* was greatly increased directly over the spots of bacteria that were grown on MEA. This did not occur with our other *Methylobacterium* sp. isolate P3, which was isolated from North Antelope Valley Creek in Lincoln, NE. It indicated that P3 is a potential antagonist against algal growth; as algae could be competition in water ecosystems as opposed to in a biological soil crust where they would be the main source of carbon. Which specific growth hormones our *Methylobacterium* spp. are producing were not identified though. It will be something we will look into in the future. The importance of Pinky’s ability (and possibly Crusty’s) (Fu et al., 2015; Hoffman et al., 2013) to produce growth-stimulating hormones will be a topic of future study as we continue to dissect the interactions involved in biological soil crust symbioses.

### Crusty has Methylobacterium spp. Symbionts Which are Likely Endosymbionts

Confirmation of Crusty’s bacterial symbionts Light Pinky and Dark Pinky have been shown through multiple experimental means. We have replicated multiple forms of fungal-bacterial endosymbiont confirming methodologies to ensure that our hypothesis of these *Methylobacterium* spp. symbiotically interacting with Crusty are accurate. For one, it seems so far that we cannot optimally grow Crusty without an actively growing Pinky. When antibiotics which kill or halt the growth of Pinky were used on Crusty, it also halted the growth of Crusty while not halting the growth of *S. cerevisiae* at the same dosage. This observation was seen in the usage of both Tetracycline and Gentamycin for all three organisms.

We replicated the bacterial live-dead staining implemented by Partida-Martinez & Hertweck to show bacterial presence within the cell wall of Crusty (2005). This result, however, can easily be contested even in the original publication, as SYTO9, the “live stain” portion is meant to stain nucleic acids. Since fungi’s nuclei and mitochondria also contain nucleic acids, it cannot be confirmed that the nucleic acids being stained are bacterial in origin. This was later acknowledged by ThermoFisher on their website after the study was performed (“SYTO™ 9 Green Fluorescent Nucleic Acid Stain,”). However, unless the nucleic acids of Crusty are spread out across the entire cell instead of in the nucleus, what we observed in these images is still likely bacterial or mitochondrial in origin. Optimally, we would like to perform immunogold labeling TEM on Crusty to confirm the presence of Pinky inside of the fungal cells. Unfortunately, this is a lengthy and expensive process and was not feasible at the time.

To molecularly determine if Pinky was an endosymbiont of Crusty or if Pinky was on the outside of the cell wall, we reproduced methods by Mondo et al., utilizing a cell surface sterilization technique to ensure only DNA that is inside of the fungal cell wall was being extracted and amplified (Mondo et al., 2012). This Chloramine-T treatment resulted in 16S rDNA amplification after treatment of Crusty’s cells, which strongly indicates that the bacterial DNA and therefore the bacteria were inside the fungal cell wall and endosymbiotic.

Results collected from the +/- carbon, nitrogen, light experiment, and the addition of antibiotics experiment, tested Crusty’s ability to interact with Pinky. If Crusty was unable to interact with Pinky, then the differences observed between the conditions in these two experiments would presumably be non-significant. However, both experiments showed that there is a significant difference in active metabolism when light is exposed to the Crusty-Pinky symbiosis in the absence of vital nutrients (p < 0.05, n=3), and that Crusty is significantly stunted in its growth in the presence of antibiotics that also stop the growth of Pinky (p < 0.01, n=3). Therefore, we can reject the null hypothesis that there are no interactions between Crusty and Pinky; however, with confidence we cannot completely confirm the hypothesis that Pinky is Crusty’s endosymbiont, only that some form of interaction is occurring.

The results displayed here show that Crusty is capable of at least bacterial symbiosis if not full endosymbiosis, which has not previously been described within polyextremotolerant fungi. While other fungal groups have been identified to have endosymbiotic bacteria (Lumini et al., 2007; Mondo et al., 2012; Partida-Martinez & Hertweck, 2005), none has yet been found in the Chaetothyriales or Dothidiales where polyextremotolerant fungi reside. Further research will need to be performed on these fungi to determine if others also harbor bacterial symbionts, if symbiosis is specific to *Methylobacterium* spp., or if Crusty is a unique outlier. However, multiple other polyextremotolerant fungi in my collection also harbor *Methylobacterium* associates, and a quick 16S rDNA amplification revealed many other fungi in my collection with positive amplification (data not shown). Indicating that Crusty is likely not the only polyextremotolerant fungus to have a *Methylobacterium* symbiont.

### Interactions of Crusty with its Methylobacterium Symbionts

We showed here that Crusty is capable of engaging in direct interactions with its bacterial symbionts, resulting in morphological changes of Crusty, increased active metabolism in nutrient-deplete conditions, and restricted fungal growth with simultaneous bacterial growth restriction in the presence of antibiotics. These observations are crucial for further understanding how the microbes in biological soil crusts are interacting for survival, and how rudimentary biofilms evolved into developing close interactions such as those seen in lichens. Additionally, specific interactions that are shown to be occurring between Crusty and its *Methylobacterium* symbionts seem to align well with other bacterial-eukaryotic symbioses.

Unique cell morphologies of the Crusty-Pinky symbiosis resemble distinct plant-bacterial and fungal-nematode interactive cell morphogenesis, particularly the root nodules that form in legume-Rhizobium symbioses (Cullimore et al., 2001; Esseling et al., 2003, 2004) and the fungal loop traps found in *Arthrobotrys oligospora* to trap nematodes (Niu & Zhang, 2011; Vidal-Diez de Ulzurrun & Hsueh, 2018). Similar morphologies in Crusty would indicate that these curved cells form in order to inoculate a fungal cell with the bacterial symbiont. Cell directionality in fungi that is not straight polar growth is generally caused by a form of chemotropism, luring the cells to grow in a specific direction that is perceived as beneficial chemical to the cell (Clark-Cotton et al., 2022; Harris, 2006; Yu et al., 2021). It has also been shown that some *Methylobacterium* spp. are capable of soybean root nodulation, implying that a similar cellular morphogenesis could be performed for fungi as well (Jourand et al., 2004; Sy et al., 2001). Without a Crusty cured of Pinky, testing this hypothesis for chemotropism and re-endosymbiosis caused by the bacteria will be difficult. However, the similarities indicates that similar mechanisms involving aforementioned symbioses and cell morphogenesis may be occurring within Crusty.

Metabolic interactions between Crusty and Pinky were of particular interest while attempting to decipher their symbiosis. We were expecting that the *Methylobacterium* spp. would contain the capabilities to perform nitrogen fixation, which is common amongst Rhizobiales (Carvalho et al., 2010). However, neither Light Pinky nor Dark Pinky contains the essential genes required for any of the three forms of nitrogen fixation (Bellenger et al., 2020; Chatterjee et al., 1997; Joerger et al., 1988). In annotating the nitrogen-fixation genes we did find the genes essential for production of bacteriochlorophyll, which are indicative of aerobic anoxygenic photosynthesis, which are evolutionarily related to the nitrogen fixing genes to the point of significant homologous protein sequences between the two separate systems (Fujita & Bauer, 2000). Other *Methylobacterium* spp. have been shown to contain genes for aerobic anoxygenic photosynthesis as well and are implied to be able to perform this function, but no direct measurement of photosynthesis has been performed (Atamna-Ismaeel et al., 2012; Zervas et al., 2019).

Results from our Crusty-Pinky culture grown in the presence or absence of light, and with or without a source of carbon or nitrogen, showed a significant increase in active metabolism when the cells were exposed to light even in the absence of carbon or nitrogen. These results indicate that a form of light-dependent metabolism must be occurring. Currently, our only known source of light-dependent metabolism in this system would be the *Methylobacterium*’s aerobic anoxygenic photosynthesis. However, it has been previously noted that bacteriochlorophyll is easily degraded when exposed to even fluorescent lights (Raser et al., 1992), which is what was used in this experiment.

If these *Methylobacterium* spp. are actually capable of performing anoxygenic photosynthesis, they are most likely to be using the bacteriochlorophyll *a* or *b* found in the *Rhodobacter* spp. (Permentier et al., 2001; Sauer et al., 1966). These bacteriochlorophylls are unique in that they absorb light at a much higher wavelength than their green relatives; they absorb light in the red to near infrared part of the spectrum (700-850 nm), a range which includes wavelengths above that which melanin has been shown to absorb (Meredith & Sarna, 2006; Tran et al., 2006). This means that when the *Methylobacterium* cells are within the melanized fungal cell wall, the melanin is not capable of blocking the wavelengths of light required for bacteriochlorophyll excitation. It has also been shown that UV light can disrupt bacteriochlorophyll *a* completely (Raser et al., 1992), giving credence to the protective inner sanctum of the melanized fungal cell.

This combination of melanin blocking UV light but allowing near infrared light, allows the *Methylobacterium* to perform aerobic anoxygenic photosynthesis while being constantly exposed to extreme UV light, if they are inside of the melanized fungal cell wall. It has also been shown, that biological soil crust growth significantly increased when grown under the exposure of only red and near infrared light (Tang et al., 2018). This light exposure type specifically enhanced for the growth of aerobic anoxygenic photosynthetic bacteria, with *Methylobacterium* amongst one of the most abundant genera, and an increase in fungal taxa (Tang et al., 2018). The UV blocking capabilities of melanin overlapping with the near infrared requiring *Methylobacterium*, could be the basis for the Crusty-Pinky endosymbiosis, but it will require more direct experiments for confirmation.

### Why is a Polyextremotolerant Fungi Harboring a Growth-Stimulating Bacteria, and How It can Help the Greater Biological Soil Crust Consortium

Ascertaining the reason for the Crusty-Pinky symbiosis is still under investigation. Whether this attempt at a simplified version of a biological soil crust was successful or not is also being analyzed. Although we still believe the essential niches for a biological soil crust remain the same (nitrogen producer, carbon producer, and biological scaffolding and/or abiotic resistance inducer), we cannot confirm that we were able to represent all three niches in the overall scheme of these experiments. For one, we can confirm that neither Light Pinky nor Dark Pinky contain the ability to perform nitrogen fixation. However, we do believe that these *Methylobacterium* spp. are performing aerobic anoxygenic bacterial photosynthesis due to the increase in active metabolism only in the presence of a light source, with the absence of vital carbon and nitrogen sources. Additionally, we have shown that Light and Dark Pinky are capable of growth stimulation of algae, which in itself could be an essential niche we had not anticipated, especially when the optimal growth periods of biological soil crusts are particularly short each year (Belnap et al., 2001; Belnap & Eldridge, 2001; Harper & Marble, 1988; Johansen, 1993; Raggio et al., 2017).

Growth stimulation of algae, cyanobacteria, and fungi, caused by *Methylobacterium*, could coincide with the specific conditions that allow for active microbial growth within the biological soil crust, increasing what could be a much slower process otherwise. We showed that in specific circumstances the bacteria will leave the confides of the fungal cells, with the most important abiotic factor influencing this escape being light availability. Since biological soil crusts are most active during the wet seasons (Harper & Marble, 1988; Miralles et al., 2012; Swenson et al., 2018), it could be that these conditions also allow Pinky to leave the fungal cells and release growth promoting hormones into the microbial biofilm. When conditions become inhospitable again, such as a lack of light in the winter months of Canada, this would reduce photosynthesis and therefore reduce the external carbon pool. Under these conditions, the *Methylobacterium* would either need to re-enter the fungus through a yet unknown mechanism that we have only observed via Crusty’s chemotropic curved cell morphology. Alternatively, there could be a maintained population of *Methylobacterium* inside of the fungal cells, which can be maintained internally to be released again when conditions improve.

If these *Methylobacterium* spp. are able to freely leave and enter the fungal cells during certain conditions/times of year, they could be surviving inside of the fungal cells by performing anoxygenic photosynthesis inside the fungi during the winter months while being free-living heterotrophic bacteria (with growth stimulating abilities) during the summer months. This would allow them the flexibility of optimizing these harsh conditions for their benefit, and their subsequent survival allows for production of growth-promoting hormones which in turn benefit the entire biological soil crust community. This ecological niche which is only possible through the melanized cell wall of Crusty and other polyextremotolerant fungi, could be an important niche in allowing the entire biological soil crust to more rapidly grow during the short optimal windows of growth conditions. Without this ecological niche, it is possible that the biological crust consortium would not be as effective in taking advantage of the short growth windows, and therefore not be as successful of a biofilm overall. Further tests will need to be performed to determine how vital the production of auxins and cytokinins are on the biological soil crust, and if the Crusty-Pinky endosymbiosis is the keystone niche for phytohormone production.

## Acknowledgements

We would like to acknowledge Tania Kurbessoian for her contribution to this manuscript in producing the ITS phylogenetic tree for Crusty and its relatives. We would also like to acknowledge Nancy M. Nguyen, who performed the algal overlay experiments as an undergraduate researcher in the Riekhof lab here at UNL.

